# How reliable is the use of ovarian dilatations for mosquito age-grading? A blinded multi-rater validation of the Polovodova method in *Anopheles* mosquitoes

**DOI:** 10.64898/2026.07.15.738643

**Authors:** Doreen Josen Siria, Halfan Ngowo, Emmanuel Hape, Mohamed Jumanne, Gerald Tamayamali, Laurent Mtima, Heather M. Ferguson, Fredros Okumu, Francesco Baldini

**Affiliations:** Environmental Health and Ecological Science Department (EHES), Ifakara Health Institute (IHI), Ifakara, Tanzania; School of Biodiversity, One Health and Veterinary Medicine (SBOHVM), University of Glasgow, Glasgow, UK; Institute of Disease Modelling, Gates Foundation, Seattle, USA; Wits Research Institute for Malaria, Faculty of Health Sciences, University of the Witwatersrand, Johannesburg, South Africa

**Keywords:** Polovodova, accuracy, precision, gonotrophic cycle, parity, nulliparous, parous

## Abstract

**Background:** Accurate estimation of mosquito age is critical for malaria surveillance, as only older female *Anopheles* mosquitoes can transmit the parasite. Traditional age-grading methods, include the Polovodova technique in which dilatations on the ovarian pedicel to tally completed gonotrophic cycles (GC) are counted. Despite their widespread use, these techniques remain poorly validated under realistic operational conditions, with limited evidence on their precision and accuracy across different parity rates, GC, and species. In this study, the accuracy and precision of the Polovodova method was evaluated across multiple age-graders and species under both laboratory and field conditions.

**Methods:** A blinded validation study was conducted using laboratory-reared *Anopheles arabiensis* and *Anopheles funestus* (N > 3,600) spanning four gonotrophic cycles, and field-collected mosquitoes from Ulanga district, Tanzania (N = 600). Three blinded researchers performed dissections and readings, while a fourth unblinded researcher served as the reference reader (R0) to obtain ground truth (for parity) or assumed truth (for GC) for laboratory mosquitoes. Accuracy and precision for parity and gonotrophic cycle classification were assessed using pairwise inter-rater comparisons, percent agreement, Fleiss’ kappa and Cohen’s kappa (κ) statistics. For field mosquitoes, which had no age-referenced group, only precision was assessed.

**Findings:** The Polovodova method accurately classified parity in both species, correctly identifying 98-100% of nulliparous and 94-96% of parous mosquitoes compared with the reference dataset. Inter-rater reliability for parity was almost perfect in *An. arabiensis* and *An. funestus* (κ = 0.81-0.83), with 82-96% pairwise agreement. GC classification was highly accurate for early stages (GC0-GC2: 85-100%) but declined at later stages GC4(34-46%), reflecting systematic underestimation of mosquito age. Overall GC reliability was moderate (κ = 0.56-0.59). Field classifications showed almost perfect agreement (κ = 0.87-0.88), supporting operational applicability.

**Conclusions:** The Polovodova method provides robust estimates of parity and early gonotrophic-cycle status in *Anopheles* mosquitoes, but late-cycle classifications are more prone to reader disagreement and systematic underestimation. Operational use should therefore rely on trained multiple-reader workflows, including adjudication of discordant or late-cycle specimens, rather than single-reader classification. These findings provide empirical benchmarks for quality-assured Polovodova age-grading in malaria vector surveillance and intervention evaluation.

## Introduction

Mosquito longevity is one of the most influential determinants of pathogen transmission (Beier et al. 1990). The ability of a female *Anopheles* mosquito to survive long enough for *Plasmodium* parasites to complete their extrinsic incubation period (EIP) directly determines whether transmission can occur, making mosquito survival a key parameter in the basic reproductive rate of malaria (Ross 1911, Macdonald G 1957, Smith et al. 2012). Early foundational work by Sir Ronald Ross introduced mathematical concepts of “pathometry” to describe malaria transmission (Ross 1911) which were later refined by Garrett-Jones through the concept of vectorial capacity, integrating entomological parameters such as mosquito density, biting frequency, and survival (Garrett-Jones 1964, Garrett Jones 1975). Daily mosquito survivorship is the most influential parameter for malaria transmission as small changes in mortality rates can significantly reduce the proportion of mosquitoes that live long enough to transmit the parasite (Smith et al. 2012). George Macdonald later expanded through sensitivity analyses the Ross-Macdonald model (Macdonald G 1956, 1957). This historical foundation established mosquito age as central to malaria epidemiology and continues to inform current surveillance efforts. Accurate age-grading methods are therefore essential for assessing mosquito longevity, evaluating vector control strategies, and understanding transmission dynamics (Detinova 1962, Gillies and Wilkes 1965). However, despite decades of research, determining the age of wild-caught mosquitoes remains challenging and often unreliable due to limitations in existing methods (Silver 2008, Brady et al. 2016).

Today, age-grading of *Anopheles* mosquitoes is done mainly via ovary dissections using either the Detinova method, which examines if ovarian tracheoles are tightly coiled indicating a nulliparous female (those that have never laid eggs) or uncoiled after egg-laying (parous female) (Detinova 1945, 1962), and/or the Polovodova method, which counts distinct dilatations on the ovarian pedicel to tally completed gonotrophic cycles (Beklemishev et al. 1959). The average gonotrophic cycle (GC) of African malaria vectors (period between taking a blood meal, laying eggs and returning to feed) has been estimated to take between 2-4 days (Fereda 2022), with variation occurring between species and in response to environmental conditions (Takken et al. 2024). *Anopheline* mosquitoes are, in general, gonotrophically concordant with each blood meal giving rise to a batch of eggs. If the length of the GC in a particular location is known and simplifying assumptions are made that mating and blood-feeding opportunities are not limited, then the age of a female mosquito can be approximated by the number of GC that have been completed.

Early work showed that each gonotrophic cycle leaves dilatations on the ovariole that can be observed under a microscope through mosquito dissection (Beklemishev et al. 1959, Anagonou et al. 2015). Entomologists can therefore apply the Polovodova method to estimate the number of gonotrophic cycles and, approximate the age of the female mosquitoes (Hugo et al. 2008, Anagonou et al. 2015). The technique involves the removal of the ovarian sheath through dissection, spreading out and separating the mosquito ovarioles and counting the number of “follicular dilatations” found in the mosquitoes ovarioles, each corresponding to a completed GC (Beklemishev et al. 1959). The physiological age of female mosquitoes is then estimated from the number of follicular dilatations, based on the assumption that each corresponds to a completed gonotrophic cycle of known duration (Beklemishev et al. 1959). Using this method, younger and older mosquitoes can be identified based on egg-laying status. However, the reliability of the method is limited by an important biological constraint: when a follicle fully develops and is ovulated as an egg, the dilatation record for that gonotrophic cycle is lost from that ovariole. It is therefore only those follicles that never fully develop typically dwarf ovarioles that retain the complete record of all past gonotrophic cycles (Hoc and Charlwood 1990, Hoc and Wilkes 1995). As a result, the proportion of ovarioles displaying the correct diagnostic number of dilatations decreases progressively with each successive gonotrophic cycle, leading to systematic underestimation of age in older, multiparous mosquitoes a limitation that has been confirmed empirically in multiple mosquito species (Cook et al. 2007, Hugo et al. 2008, Anagonou et al. 2015).

While both the Detinova and Polovodova method can be used estimate mosquito age based on reproductive status, the Detinova method is limited to binary classification (nulliparous vs. parous), providing no information on the number of completed gonotrophic cycles (Hugo et al. 2008). Thus, although simpler, quicker and more widely used than the Polovodova method, the Detinova approach is not suitable for fine-scale description of mosquito age structure (Detinova 1945, Gillies and Wilkes 1965, Johnson et al. 2020). While the Polovodova can provide this higher resolution age-grading, it has the disadvantage of requiring significantly greater skill since even slight damage to the ovaries during dissection can obscure the dilatations, making age estimation difficult (Detinova 1962). This method is challenging to perform on *Anopheles* species, due to their relatively small body size, leading to frequent misinterpretation of results (Pretorius et al. 2023). It is also time-consuming and demands significant expertise (Gillies 1954, Detinova 1962, Hugo et al. 2008).

Although newer techniques for mosquito age estimation are emerging, these two ovary-dissection methods remain widely used in vector control despite notable limitations. Both methods require freshly killed mosquitoes dissected immediately after collection (Hugo et al. 2008, Anagonou et al. 2015). Although earlier studies have evaluated these methods, important gaps remain in its standardization across species, readers and gonotrophic stages, limiting cross-study comparability and routine operational use. Unlike the Detinova binary parity classifications, the Polovodova method has greater potential for fine-scale GC classification, yet its accuracy and precision have rarely been formally evaluated under controlled conditions for major malaria vectors.

A critical consideration in evaluating mosquito age-grading methods is distinguishing between individual-level classification accuracy and the consistency of population-level age structure estimates. While accurate individual GC classification is biologically important, malaria surveillance programmes primarily rely on age structure data to estimate mosquito survival and assess vector control impact. Thus, a method can still provide useful surveillance information if misclassifications are consistent and reproducible across observers, even with some individual-level errors.

In this study, we therefore conducted a comprehensive evaluation of the Polovodova age-grading method in two major African malaria vectors, *Anopheles arabiensis* and *Anopheles funestus*. Specifically, we aimed to assess the accuracy and precision of i) parity classification (nulliparous vs parous) and ii) gonotrophic cycle classification of lab-reared mosquitoes with known physiological status under controlled conditions across successive blood-feeding stages; and lastly to iii) determine the consistency and reproducibility of the method using the same graders in estimating the age of wild caught *An. arabiensis* and *An. funestus*.

## Methods

### Mosquito rearing and handling

Experiments were conducted at the Ifakara Health Institute’s VectorSphere facility in Ifakara, Tanzania. Laboratory-reared *An. arabiensis* and *An. funestus* mosquitoes from established colonies maintained at the Ifakara Health Institute were used. The *An. funestus* colony originated from field collections and was recently established and maintained at IHI as described by Hape *et al*. (Hape et al. 2025), while the *An. arabiensis* colony has been maintained at IHI since the early 2010s following collections from Sagamaganga, Tanzania (Ng’Habi et al. 2010). The laboratory-reared *An. arabiensis* and *An. funestus* mosquitoes were maintained under controlled temperature, humidity, and photoperiod. Each experimental replicate consisted of separate cohorts (six replicates for *An. funestus* and seven for *An. arabiensis*).

Upon emergence, mosquitoes were maintained on 10% sucrose solution and were not offered any blood meal until the scheduled experimental feeding. Male mosquitoes were maintained alongside females throughout the experimental period to ensure natural mating within cohorts. Females were only separated at scheduled oviposition time points to allow individual egg-laying assessment. The first blood meal was provided at day 4 post-emergence, after an initial sugar-feeding period allowing for adult maturation. Mosquitoes were provided with human blood meal every 4 days, followed by oviposition opportunities two days after each blood-meal, simulating up to four gonotrophic cycles. We chose a 4-day interval between blood meals to approximate realistic gonotrophic cycle lengths previously reported for *An. funestus* and *An. arabiensis,* which span approximately 3-5 days under favourable conditions (Gillies and Wilkes 1965, Ngowo et al. 2021, Mwanga et al. 2024). Each replicate included approximately ∼300 female mosquitoes for each of the species, ensuring and allowing us to obtain enough for subsequent parity and gonotrophic cycle assessment (GC0-GC4), typically targeting <’25-30 individuals per category.

### Blood feeding and oviposition

For each species, 2,000-3,000 pupae were placed in 30 cm^2^ cages. Upon emergence, mosquitoes were provided with a 10% sucrose solution continuously. Blood meals were provided on a set schedule to initiate and track gonotrophic cycles as follows (Figure 1)

**Figure 1.**
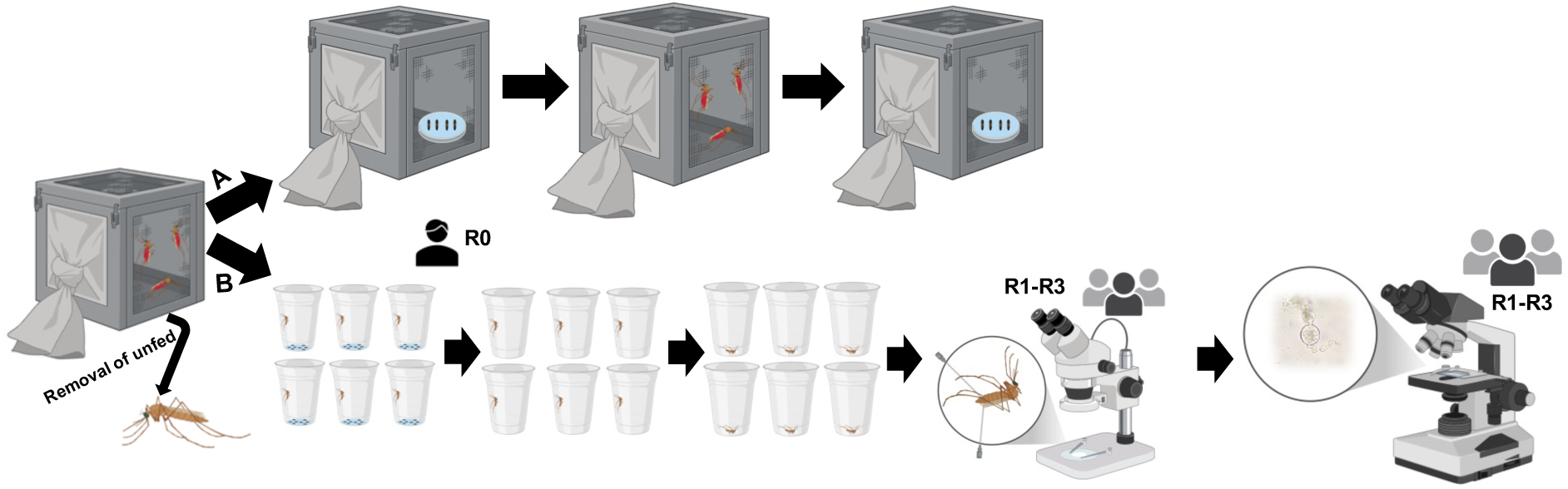
Schematic overview of the experimental design and workflow for mosquito blood-feeding, oviposition, dissection, and age-grading. First, adult female mosquitoes were blood-fed and unfed individuals removed. Then, A) oviposition cups were placed inside the cage for group oviposition and simultaneously B) 60 mosquitoes were separated for individual oviposition. Individual mosquitoes were recorded for presence/absence of laid eggs by Researcher 0 (R0), euthanized and dissected under a stereomicroscope by three researchers (R1-R3, 20 mosquitoes each), blind to the physiological states. Age-grading based on ovarian morphology was determined under a compound microscope, with slides exchanged among researchers for blinded assessment. Mosquitoes that laid eggs in the cage (A), where subsequently blood fed and procedure B repeated up to the fourth blood meal. To obtain nulliparous mosquitoes, a subset of females was collected prior to the first blood meal and directly dissected and age-graded by R1-R3.

On days 1-2 post-emergence, a subsample of 60 female mosquitoes were randomly collected from each cage prior receiving their first blood meal and were dissected to serve as the “known” nulliparous (GC0) reference group. The remaining mosquitoes in the large cages were then blood-fed on day 4 for the first time. On day 6, a new random subset of 60 females were removed from the main cage and transferred to individual oviposition cups, while the rest of the mosquitoes were allowed to oviposit communally in plastic bowls which were placed inside the cages. All mosquitoes (both individually separated and those in communal cages) had continuous access to 10% sucrose solution and oviposition substrates.

This procedure was repeated for subsequent gonotrophic cycles. Specifically, new random subsets of 60 females were collected from the main large cages for dissection after the second, third, and fourth blood meals on days 8, 12, and 16, respectively.

These subsets were therefore independent and parallel, drawn from the large source population rather than sequentially following the same individuals. After each blood-feeding event, females that were visually confirmed as unfed (no visible blood in the abdomen and no abdominal distension) were removed from the cages to reduce potential confounding.

### Linkage between blood feeding events and gonotrophic cycles

In this study, blood feeding events (BF) and gonotrophic cycles (GC) were directly linked. Each successful blood meal followed by egg-laying can advance the mosquito to the next gonotrophic cycle. Therefore, BF0 corresponds to GC0 (nulliparous, before the first blood meal), BF1 after egg laying corresponds to GC1 (after the first blood meal and oviposition), and so on. This experimental design enabled direct comparison between the known physiological history of each mosquito (based on controlled blood feeding and oviposition records) and the gonotrophic age assigned by dissectors during ovarian examination.

### Dissection and ovarian assessment of lab-reared mosquitoes

The subset of mosquitoes moved into individual holding cups were removed, they had oviposited and were then euthanized using chloroform; after which their ovaries were dissected within 10-15 minutes after killing within the same day. All individuals that laid eggs following the first blood meal could be confirmed as having completed one gonotrophic cycle. However, the gonotrophic cycle was uncertain for females observed to lay eggs on the second, third or fourth blood meal as their previous egg laying history (before the current blood meal) was not directly observed. Thus, a model was used to estimate the ‘assumed ground truth’ of GC completed by mosquitoes on later feeding rounds; based on accounting for the number of previous blood feeding opportunities and the proportion of mosquitoes that required more than one blood feed to produce eggs. The number of gonotrophic cycles was determined using the Polovodova age grading method (Beklemishev et al. 1959). Nulliparous females were defined as those with no ovarian dilatations, while parous females were identified as having one or more dilatations (graded as GC1, GC2, GC3 or GC4 based on number of dilations observed). This experimental procedure was replicated to achieve a target sample size of approximately 1,800 mosquitoes per species, per blood meal (BF0 to BF4).

The reference researcher (R0) reared and blood-fed mosquitoes under controlled conditions and had full knowledge of their age and blood feeding history, while the other three researchers (R1, R2, and R3) were blinded. For each replicate, R0 prepared 60 mosquitoes for dissection and age-grading by transferring them into uniquely labelled cups (without eggs), which were blindly and equally distributed for dissection to R1, R2, and R3 (20 mosquitoes each), who independently dissected and age-graded the female mosquitoes using the Polovodova age-grading method. Afterward, R1-R2-R3 researchers exchanged slides for additional, independent evaluations, allowing each sample to be age-graded by multiple observers. All dissections were performed under stereo microscopes, and results read using compound microscopes (Figure 2).

**Figure 2.**
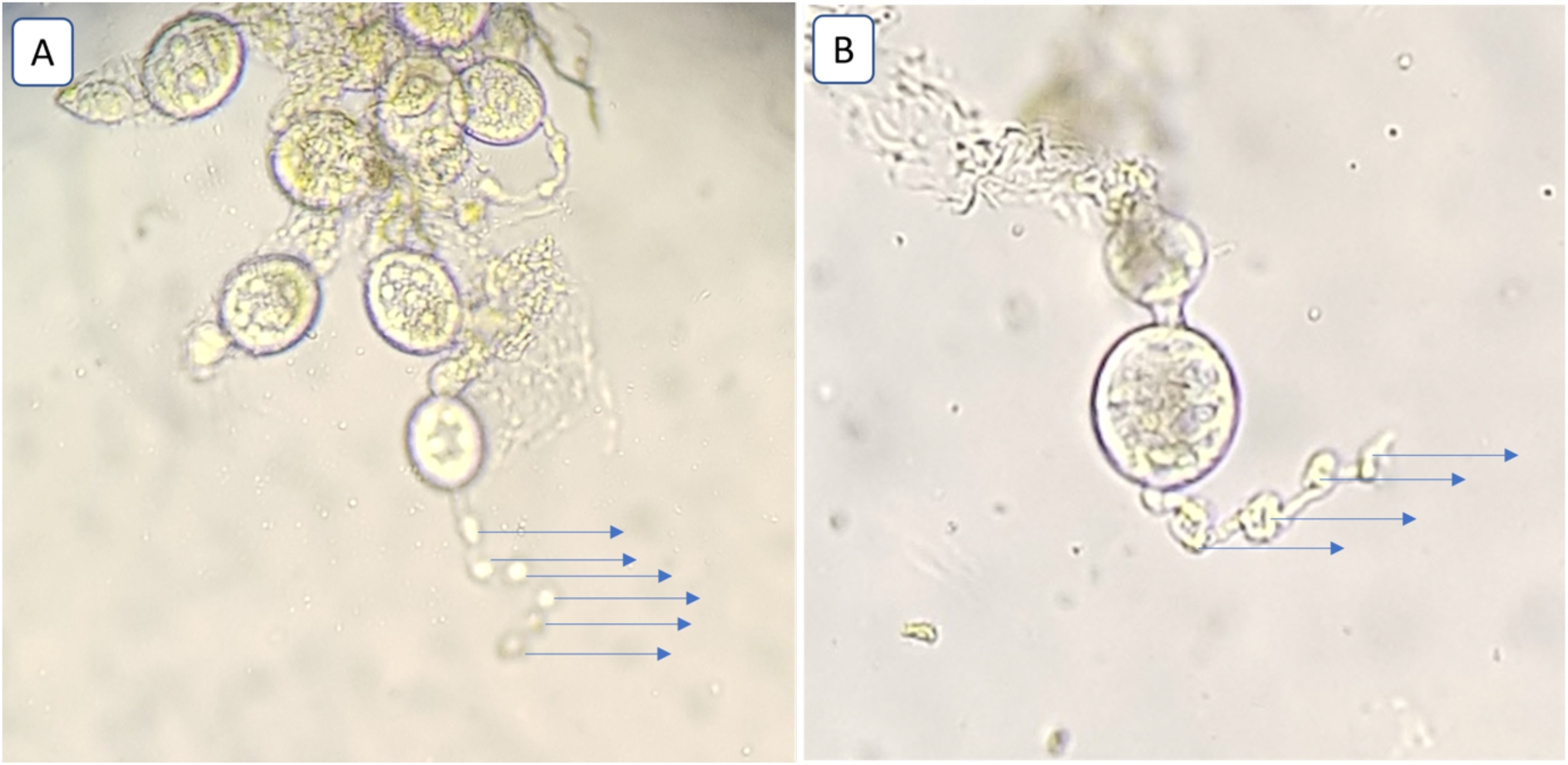
Ovarian dilatations indicating completed oviposition events. Blue arrows highlight follicular dilatations on the ovarioles corresponding to the number of oviposition performed by the mosquito: (A) six times egg oviposition and (B) three times egg oviposition.

### Establishing standard reference for parity assessment in laboratory-reared mosquitoes

The true parity status of the tested mosquitoes was established from direct observations of blood-feeding and oviposition events by the reference researcher (R0). Where oviposition was individually observed, parity status was determined with certainty based on whether a mosquito laid eggs following a blood meal: mosquitoes observed to lay eggs after their first blood meal were assigned confirmed parous status, while those that did not lay eggs were classified as nulliparous. For later gonotrophic cycles, because oviposition occurred collectively within cages, it was not possible to track the egg-laying history of individual mosquitoes. Consequently, mosquitoes that had received multiple blood meals but did not lay eggs after their most recent oviposition opportunity could not be definitively classified and were excluded from the analysis. As illustrated in the supplementary Figure S1, only mosquitoes with clearly observable oviposition outcomes following their most recent blood meal were reserved for parity analysis (Supplementary Figure S1). The dissections conducted by R1-R3 were performed independently and were subsequently compared against this reference classification to evaluate the accuracy of the Polovodova method.

### Establishment of assumed gonotrophic cycles in laboratory-reared mosquitoes

Because egg laying was directly observed only after the final blood meal, the complete reproductive history of mosquitoes given multiple blood-feeding opportunities was unknown. We therefore created an assumed reference (AR) dataset using available records from R0, including blood-feeding history and observed oviposition, together with dissection observations from R1-R3, including whether eggs were present in the abdomen. This approach assumed that each blood meal usually resulted in one oviposition event, while allowing for possible gonotrophic discordance (Lardeux et al. 2008, Derek Charlwood et al. 2016), where more than one blood meal may be required for a single egg batch (Briegel and Hörler 1993, De Oliveira et al. 2012). The decision framework used to infer assumed gonotrophic cycle categories is illustrated in the supplementary Figure S2, based on the following assumptions: 1) Mosquitoes that laid eggs after the final blood meal were assigned to the corresponding gonotrophic cycle. 2) Those that did not oviposit but had eggs in the abdomen at dissection were classified as gravid. 3) For mosquitoes that neither oviposited nor appeared gravid after the second to fourth blood meal, gonotrophic cycle status was assigned probabilistically, with 90% assigned to the preceding cycle and 10% to two cycles earlier. 4) Finally, the proportion of mosquitoes assigned to the GC corresponding to the same number of blood meals was calculated as the proportion that laid eggs minus the proportion assigned to the preceding GC. Although this AR dataset was not a true “ground truth”, it provided a structured, biologically informed reference for evaluating gonotrophic cycle classifications in the absence of complete individual oviposition histories (Supplementary Figure S2).

### Field-collected mosquitoes

To evaluate the precision of Polovodova age-grading under field conditions, we compared the ovarian-dilatation readings reported independently by four independent researchers (R1-R4). Wild mosquitoes were collected over multiple days using CDC light traps from Tulizamoyo village (8.354497°S, 36.705468°E), where *Anopheles arabiensis* predominates, and Ebuyu village (8.97900° S, 36.7600°E), where *An. funestus* is the dominant vector, both in Ulanga District, south-eastern Tanzania. A total of 600 females were independently dissected and classified as nulliparous or parous; parous females were further assigned to gonotrophic cycle categories based on the number of ovarian dilatations observed. Because the true reproductive history of field-collected mosquitoes was unknown, accuracy could not be assessed; therefore, this analysis focused on precision, measured as inter-rater agreement among researchers.

### Data analysis

Statistical analyses were performed to evaluate two key properties; (i) accuracy (concordance between the age grades assigned by readers and the known physiological age of the mosquitoes) and (ii) precision (consistency of age classification between different readers). Both accuracy and precision were assessed by quantifying inter-rater reliability (IRR) among the three graders (R1-R3).

The accuracy assessment was evaluated using the true or assumed true values established by R0. For parity classification (nulliparous vs. parous), the parity status assigned by each researcher (R1-R3) was compared with the reference classification established by R0 based on direct observations of blood feeding and egg-laying history (Figure 2). The proportion of correctly classified mosquitoes was calculated to estimate the accuracy of each researcher (R1-R3), and agreement between each researcher and R0 was additionally assessed using Cohen’s *kappa* to determine whether observed agreement exceeded chance.

For gonotrophic cycle (GC) classification, the distribution of mosquitoes across GC categories assigned by each researcher (R1-R3) was compared with the expected age structure derived from the assumed reference (AR) dataset generated from R0 egg-laying records (supplementary Figure S1). The distribution of mosquitoes assigned to each category (GC0-GC4) by the graders were compared with the expected distribution relative to AR dataset based on the number of completed blood-feeding events. Graphical comparisons and distribution plots were generated, stratified by blood-feeding event.

Agreement between observed and expected gonotrophic cycle distributions was quantified using a distributional overlap score:

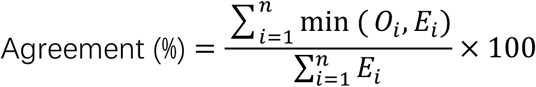

where E_i_ represents the expected count from the AR dataset in gonotrophic cycle category i, O_i_ represents the observed count assigned by the researcher in gonotrophic cycle category, i (where i can be GC0, GC1, GC2, GC3, GC4), and Σ represents summation across all five GC categories. A small constant (α=0.5) was added to address structural zeros. This metric ranges from 0 (no overlap) to 1 (perfect overlap). Ninety-five percent confidence intervals were obtained through 20,000 simulations from a Dirichlet distribution using the Highest Posterior Density (HPD) interval. All statistical analyses were performed in R software separately for *An. arabiensis* and *An. funestus* and stratified by blood-feeding event (BF0-BF4).

Precision for both parity (nulliparous vs. parous) and gonotrophic cycle (GC0-GC4) classifications was evaluated through pairwise agreement between graders using Cohen’s kappa statistic (κ), while overall agreement among all three graders was assessed using Fleiss’ kappa. These statistics were computed using the irr package in R software. Kappa values were interpreted according to the commonly used scale: ≤0 = no agreement, 0.01-0.20 = slight agreement, 0.21-0.40 = fair agreement, 0.41-0.60 = moderate agreement, 0.61-0.80 = substantial agreement, and 0.81-1.00 = almost perfect agreement (Cohen 1960). For each κ estimate, the kappa statistic is reported alongside its 95% confidence interval and p-value. A total of more than 3,600 mosquitoes from replicate experiments involving both *An. arabiensis* and *An. funestus* were included in the analyses.

For field-collected mosquitoes, only precision (inter-rater reliability) was assessed, given the absence of a known reference history. Agreement among four readers (R1-R4) was evaluated for both parity (nulliparous vs. parous) and GC classification. Gravid mosquitoes were excluded. Pairwise agreement was measured using Cohen’s kappa and overall agreement using Fleiss’ kappa. Confusion matrices were generated to visualise patterns of agreement.

## Results

### Accuracy of parity classification in laboratory-reared females

All three researchers (R1-R3) achieved high accuracy in identifying parous and nulliparous females, when compared against the reference reader, R0. For both *An. arabiensis* and *An. funestus*, accuracy was 98-100% for nulliparous and 94-96% for parous mosquitoes. Agreement between each researcher and the reference was significantly high, with Cohen’s *kappa* values of κ = 0.87-0.89 for *An. arabiensis* and κ = 0.86-0.92 for *An. funestus* (all p < 0.001, Table 2).

### Accuracy of gonotrophic cycle classifications in laboratory-reared females

The accuracy of gonotrophic cycle (GC) classification by the three researchers (R1-R3) was evaluated against the Assumed Reference (AR) dataset for *An. arabiensis* (N=420) and *An. funestus* (N=360). Results are presented in Figure 4, Figure 5 and supplementary Tables S2-S4.

**Figure 3.**
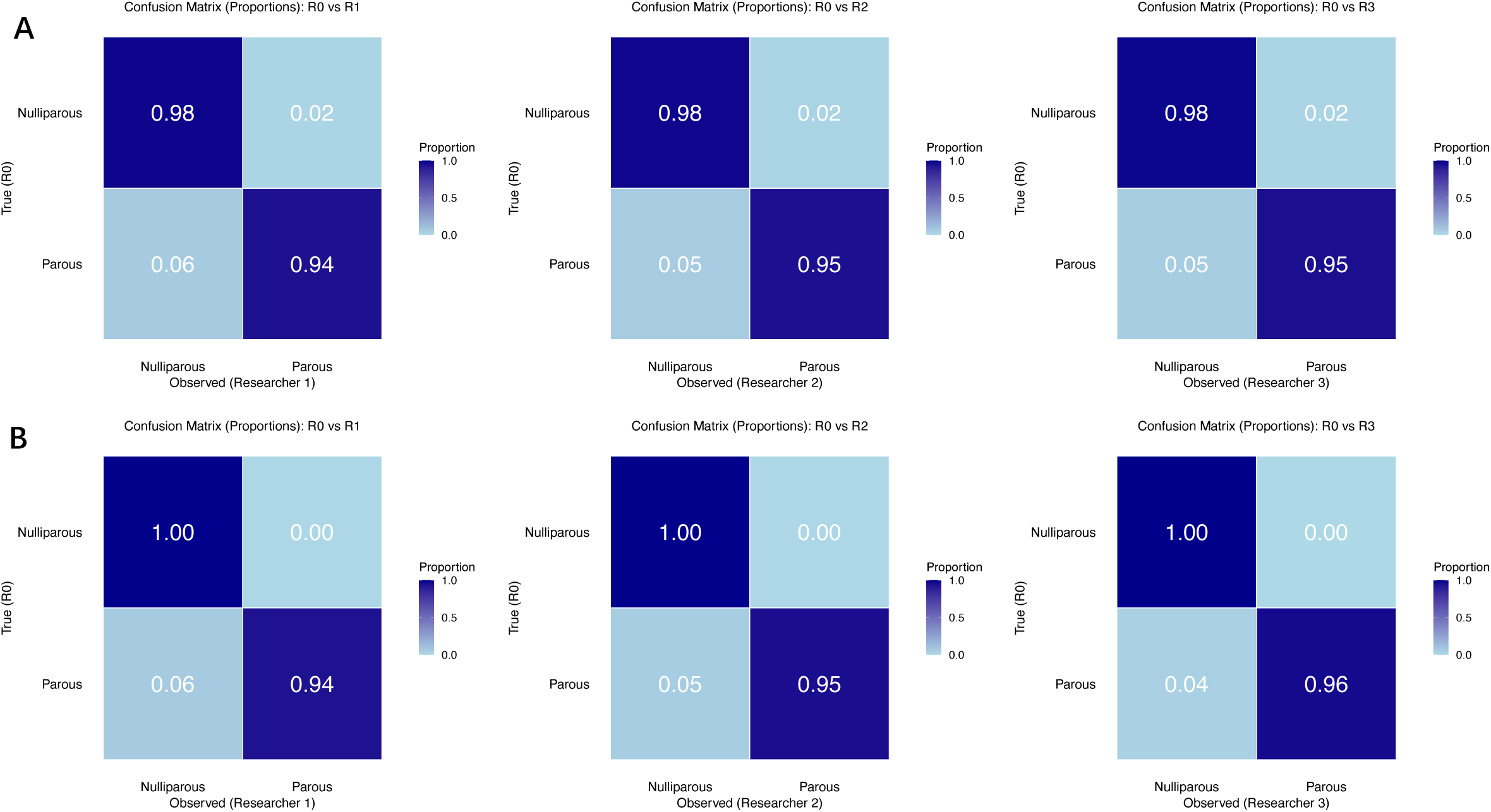
**Accuracy of parity classification.** Confusion matrices comparing parity classifications (nulliparous vs. parous) observed by researchers R1, R2, and R3 against reference data (R0) for (A) *An. arabiensis* and (B) *An. funestus*. Each row represents true parity states (assessed by R0), and columns represent the classification determined by R1, R2 and R3.

**Figure 4.**
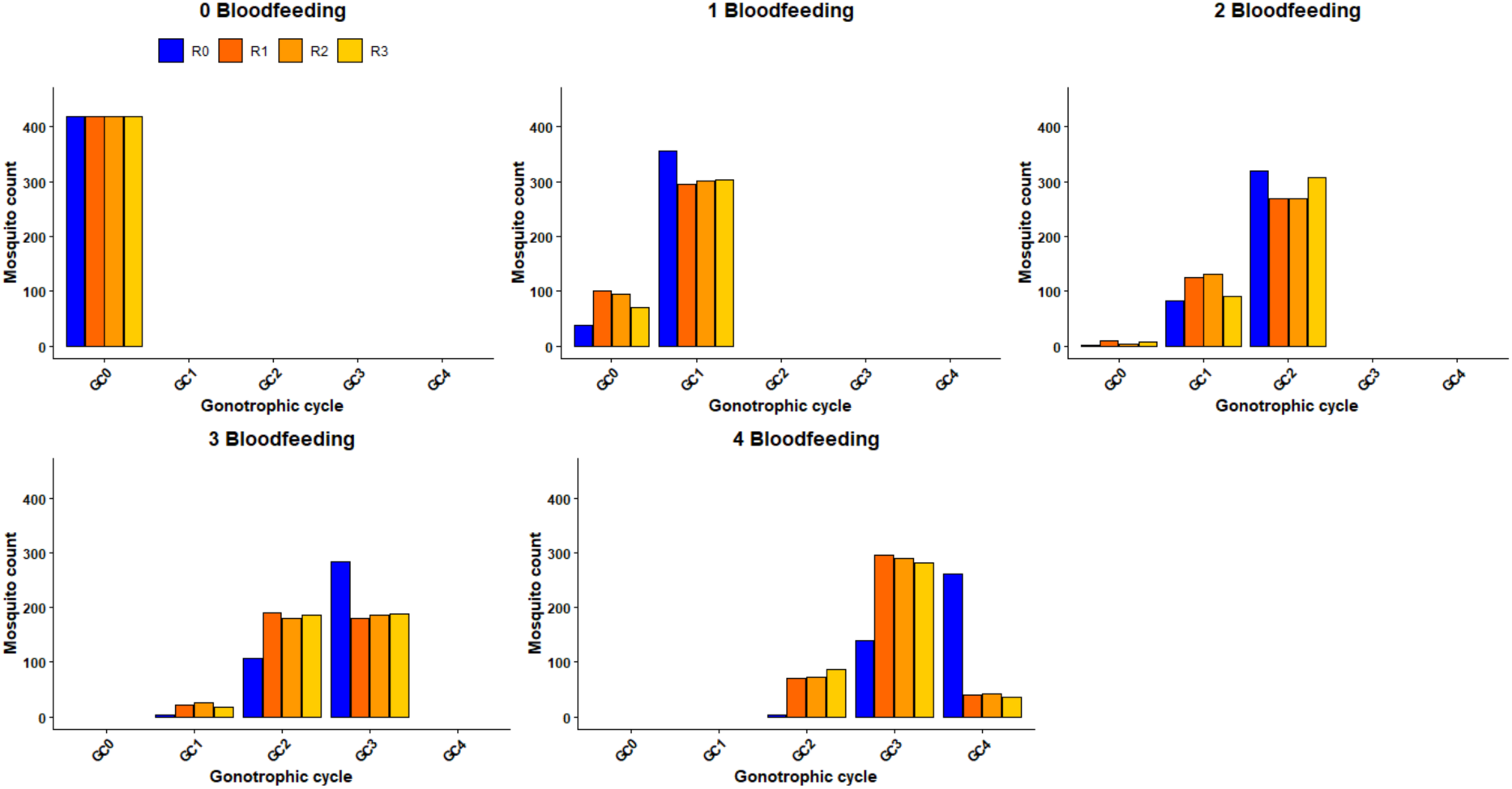
Accuracy of gonotrophic cycle classification for *An. arabiensis*. Agreement between researcher-assigned gonotrophic cycle classifications and the assumed reference (AR) age structure at different blood meals (0 to 4).

**Figure 5.**
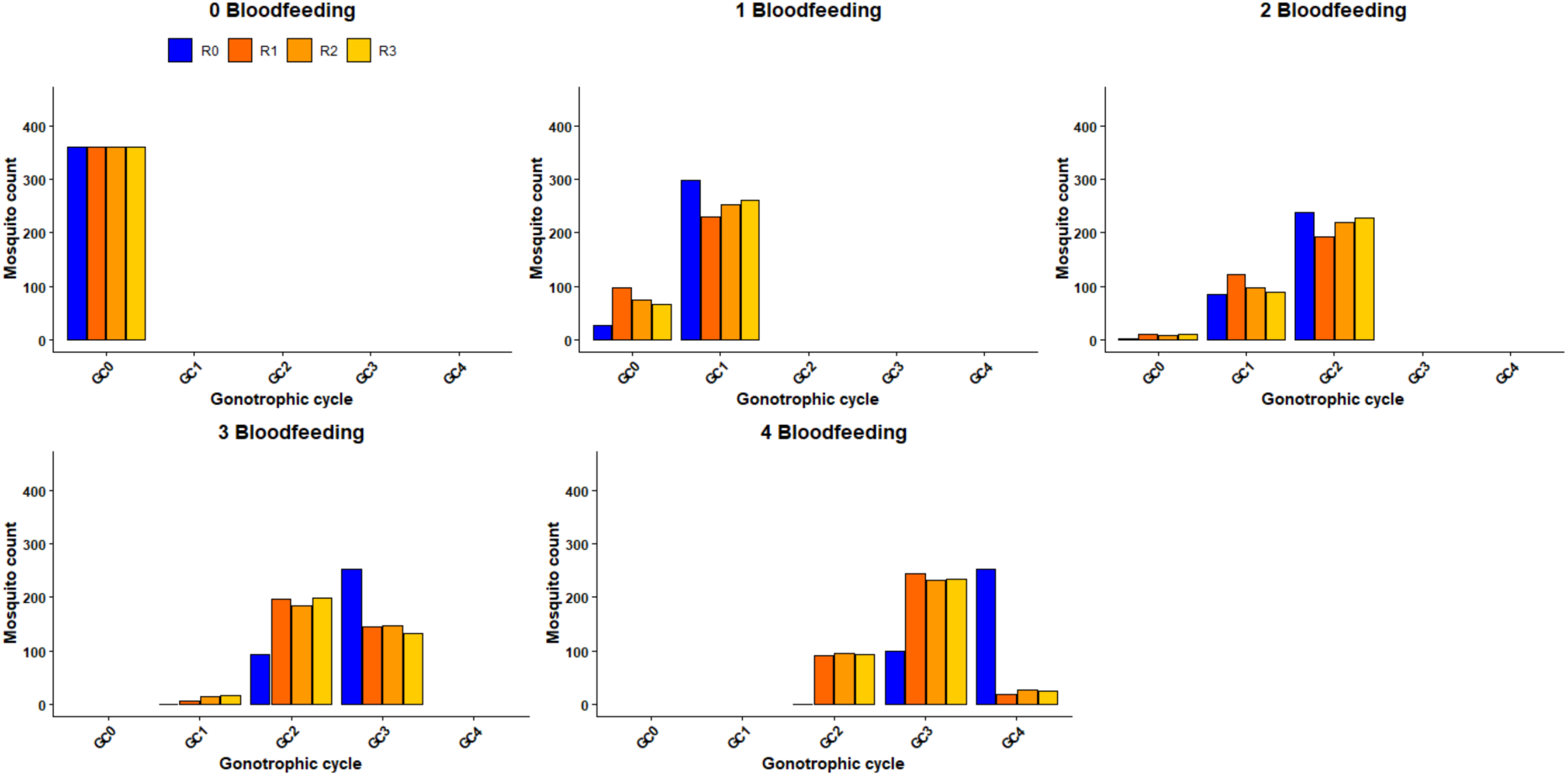
Accuracy of gonotrophic cycle classification for *An. funestus*. Agreement between researcher-assigned gonotrophic cycle classifications and the assumed reference (AR) age structure at different blood meals (0 to 4).

Visual inspection of the distributions of *An. arabiensis*, Figure 4 and of *An. funestus*, Figure 5 showed strong agreement between researcher classifications and the AR dataset for early gonotrophic cycles. All researchers achieved perfect classification (100%) for nulliparous mosquitoes (BF0). Agreement remained high after the first and second blood meals but declined noticeably in later cycles, with researchers consistently underestimating mosquitoes in GC3 and GC4.

This visual pattern was confirmed by distributional overlap analysis. For *An. arabiensis*, overlap scores were excellent at BF0 (1.00) and remained high during BF1 (0.84-0.86) and BF2 (0.88-0.97). However, agreement declined at BF3 (0.75-0.76) and dropped to poor levels at BF4 (0.45-0.46) (supplementary Table S2). A similar trend was observed in *An. funestus*, with perfect agreement at BF0, good agreement at BF1 and BF2, and poor agreement at BF3 (0.65-0.69) and BF4 (0.34-0.36) (supplementary Table S2).

The decline in accuracy at higher gonotrophic cycles was consistent across all three readers, as shown by narrow 95% confidence intervals. These findings indicate that while the method performs well for nulliparous and young parous mosquitoes, its accuracy is limited for older mosquitoes (≥3 gonotrophic cycles), likely due to increasing difficulty in distinguishing follicular dilatations (supplementary Table S3-S4).

### Precision of parity classifications in laboratory-reared females

The precision of the Polovodova method in classifying the nulliparous and parous *for An. arabiensis* among researchers R1, R2, and R3 demonstrated high agreement across all parity categories (p < 0.001). Parity classification agreement was consistently high, with 84-86% for nulliparous and 95% for parous mosquitoes. Overall inter-rater reliability across all the three researchers, estimated using Fleiss’ kappa, was 0.81 [95% CI: 0.80-0.82], indicating substantial consistency among researchers (Figure 6). Similarly, parity status for *An. funestus* was assessed with high precision among graders, with an overall Fleiss’ kappa of 0.83 [95% CI: 0.82-0.84; p < 0.001]. Agreement for nulliparous was 82%-89% agreement and for parous mosquitoes was 95%-97%. Inter rater agreement ranged from κ = 0.80 to 0.88 (Figure 6, supplementary Table S5).

**Figure 6.**
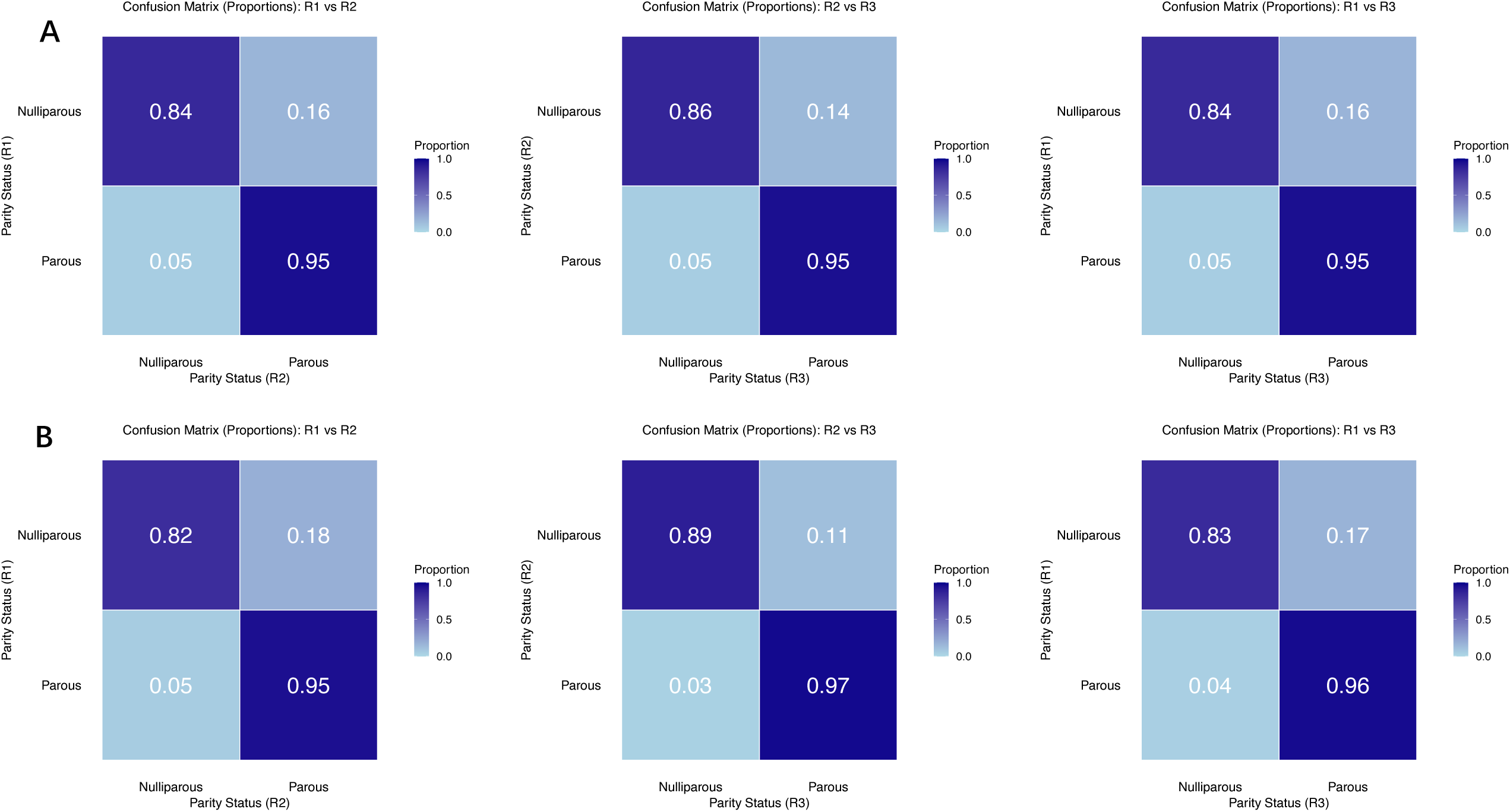
Precision of parity classification. Confusion matrices comparing pairwise agreement classification (nulliparous vs parous) between researchers (R1 vs. R2, R2 vs. R3, R1 vs. R3) for (A) *An. arabiensis* and (B) *An. funestus*. Each row and column represent the parity states determined by each researcher.

### Precision of gonotrophic cycle classifications in laboratory-reared females

The precision for *An. arabiensis* among researchers (R1 vs. R2, R2 vs. R3, and R1 vs. R3) revealed strong agreement and high consistency in identifying nulliparous mosquitoes (GC0), with agreement of 84-86% across comparisons (p < 0.001). However, precision declined for later gonotrophic cycles (GC1-GC3), with an agreement of 60-76% across comparisons. The inter-rater reliability indicated moderate agreement across researchers [κ = 0.58-0.60, 95% CI: 0.56-0.63], reflecting the observed decrease in precision for later cycles (Figure 7; Table S6). Overall, precision patterns were broadly similar between species, although estimates for *An. funestus* showed slightly lower agreement across researchers compared with *An. arabiensis.* Similarly, for *An. funestus*, pairwise comparisons demonstrated significant agreement for GC0 (82-89%) but precision declined for later gonotrophic cycles, GC1-GC3 was 59-69%. Pairwise Cohen’s kappa values indicated moderate agreement among researchers [κ = 0.55-0.58, 95% CI: 0.52-0.61], supporting the observed variability in estimating gonotrophic cycles beyond the nulliparous stage (p < 0.001; Figure 7; Table S6). Overall agreement among all three researchers was similarly moderate, as indicated by Fleiss’ kappa was [κ = 0.59, 95% CI :0.58-0.61] and [κ = 0.56, 95% CI: 0.54-0.59] for *An. arabiensis* and *An. funestus* respectively.

**Figure 7.**
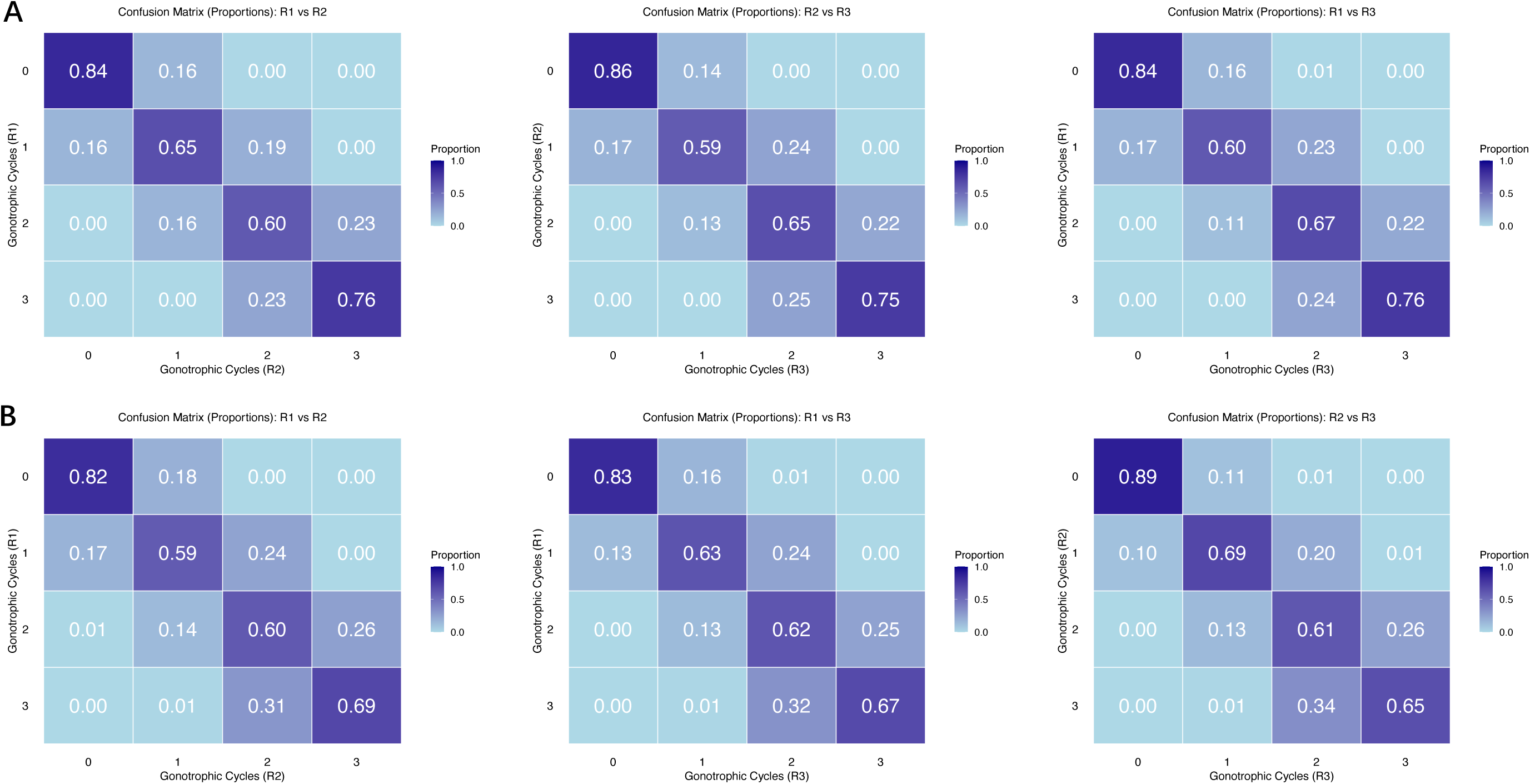
Precision of gonotrophic cycle classification. Confusion matrices showing the precision percentages of the gonotrophic cycles of (A) *An. arabiensis* and (B) *An. funestus* between researchers. Each matrix compares the gonotrophic cycle between two researchers (R1 vs. R2, R2 vs. R3, R1 vs. R3).

### Precision of parity and gonotrophic cycle classifications in field-collected mosquitoes

Parity-classification precision was assessed in 559 field-collected *An. arabiensis* and 507 *An. funestus*, while gonotrophic-cycle precision was assessed in 521 and 459 mosquitoes, respectively. During dissections, mosquitoes spanning multiple gonotrophic stages were identified, including GC0 (nulliparous), parous with sac (recently oviposited parous females, collected within 24 hours after egg laying and identified by a visible ovarian sac), GC1 (mosquitoes that had completed one gonotrophic cycle), and GC2 (mosquitoes with two completed gonotrophic cycles). Because the recently oviposited and GC2 categories each contained fewer than 20 mosquitoes per species, they were excluded to avoid unstable estimates, and gonotrophic-cycle precision analyses restricted to GC0 and GC1.

#### Parity and gonotrophic cycle classifications for field-colected An. arabiensis

For *An. arabiensis*, overall agreement in parity classification among the four researchers was significantly high (Fleiss’ κ = 0.87, p < 0.001). Pairwise Cohen’s kappa values ranged from κ = 0.82 (95% CI: 0.76-0.88) to κ = 0.933 (95% CI: 0.89-0.97), indicating consistently high agreement across all researcher pairs (p < 0.001). Pairwise confusion matrices showed strong concordance for both nulliparous (85-94%) and parous classifications (96-99%), Figure 8A, Table S7). Similarly, overall agreement in GC classification among the four researchers was very high (Fleiss’ κ = 0.88, p < 0.001). Pairwise Cohen’s kappa values were between κ = 0.83 (95% CI: 0.77-0.89) and κ = 0.93 (95% CI: 0.89-0.97), demonstrating strong agreement between researchers (p < 0.001). Agreement was 86%-94% for GC0 and 96%-98% for GC1 (Figure 8B, Table S8).

**Figure 8.**
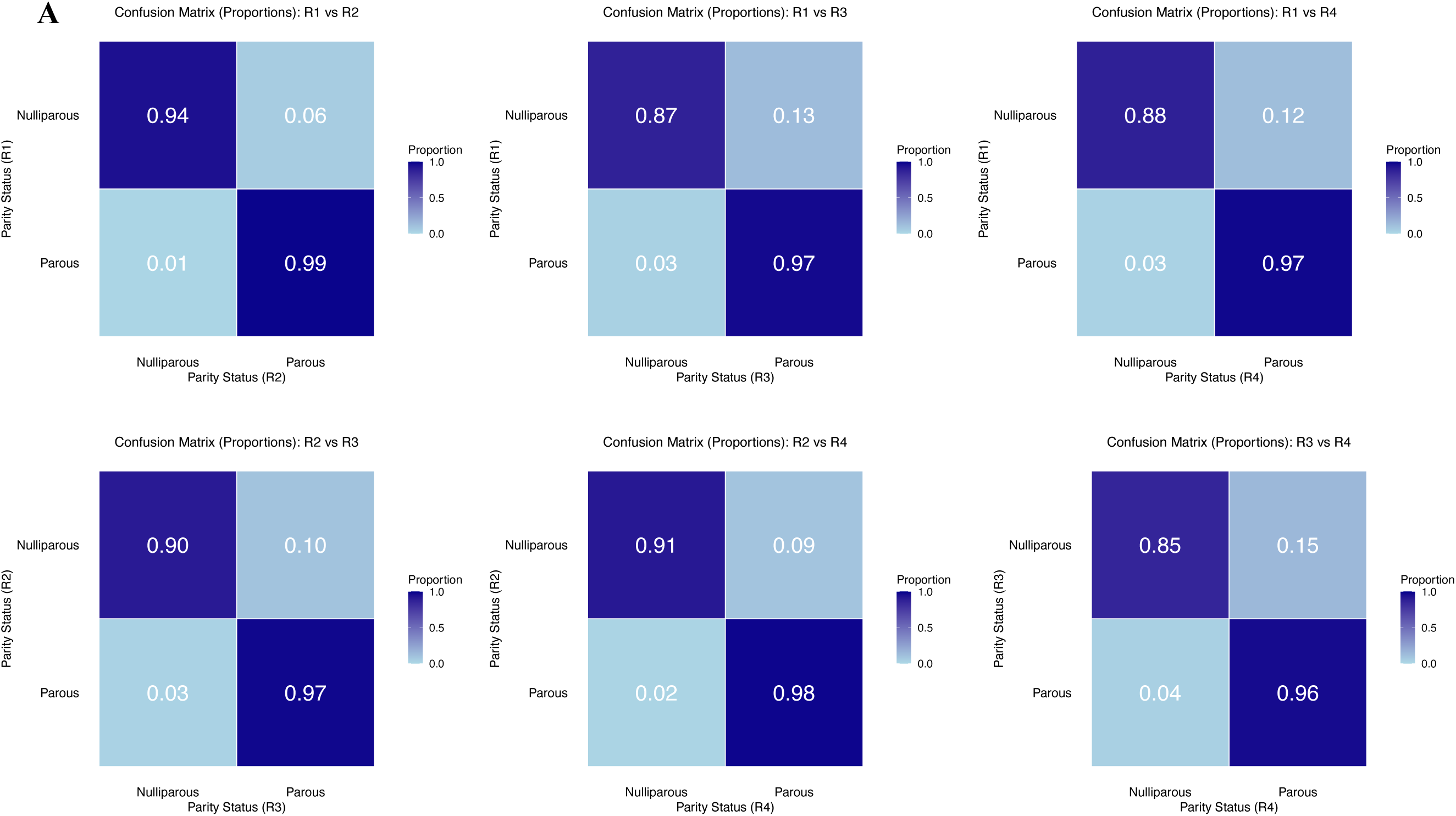
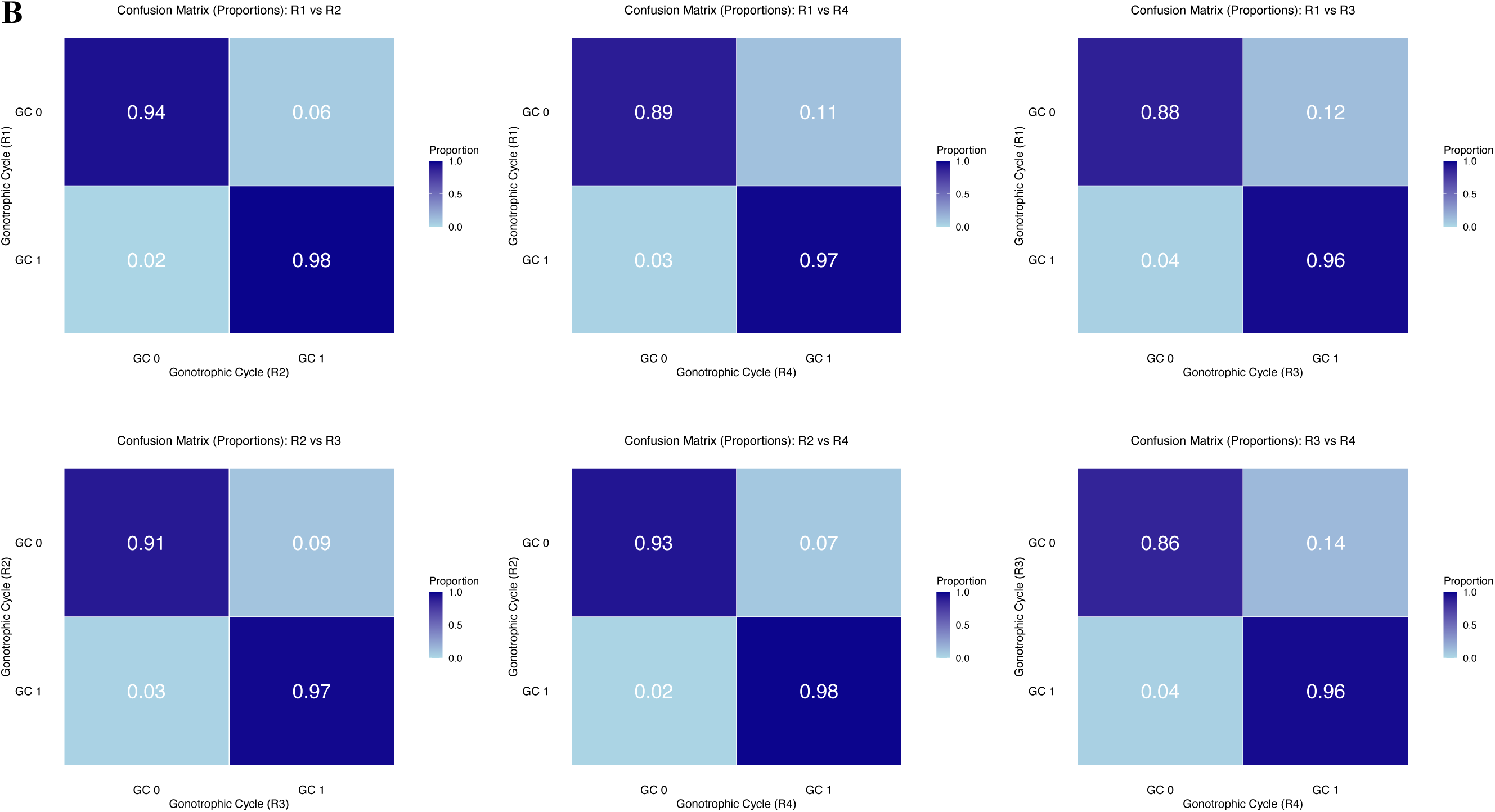
Precision of A) parity and B) gonotrophic cycle classification in field-collected *An. arabiensis*. Confusion matrices showing pairwise agreement in parity (nulliparous vs parous) and gonotrophic cycle classification (GC0 and GC1) between researchers (R1 vs R2, R1 vs R3, R1 vs R4, R2 vs R3, R2 vs R4, and R3 vs R4).

#### Parity and gonotrophic cycle classifications for field-collected An. funestus

For *An. funestus*, overall agreement in parity classification among the four researchers was high (Fleiss’ κ = 0.861, p < 0.001), with pairwise Cohen’s kappa values between κ = 0.79 (95% CI: 0.72-0.85) and κ = 0.93 (95% CI: 0.89-0.97), indicating strong agreement across all researcher pairs (p < 0.001). Agreement for identifying nulliparous was (85%-96%) and 94%-99% for parous mosquitoes, (Figure 9A, Table S7). Similarly, the overall GC classification agreement was high with Fleiss’ κ of 0.87 (p < 0.001) with pairwise Cohen’s kappa values between κ = 0.79 (95% CI: 0.73-0.86) and κ = 0.93 (95% CI: 0.89-0.97), reflecting strong consistency in GC classification among researchers (p < 0.001). High agreement was observed for both GC0 (86-96%) and for GC1 was (94-99%) across researchers, (Figure 9B, Table S8).

**Figure 9.**
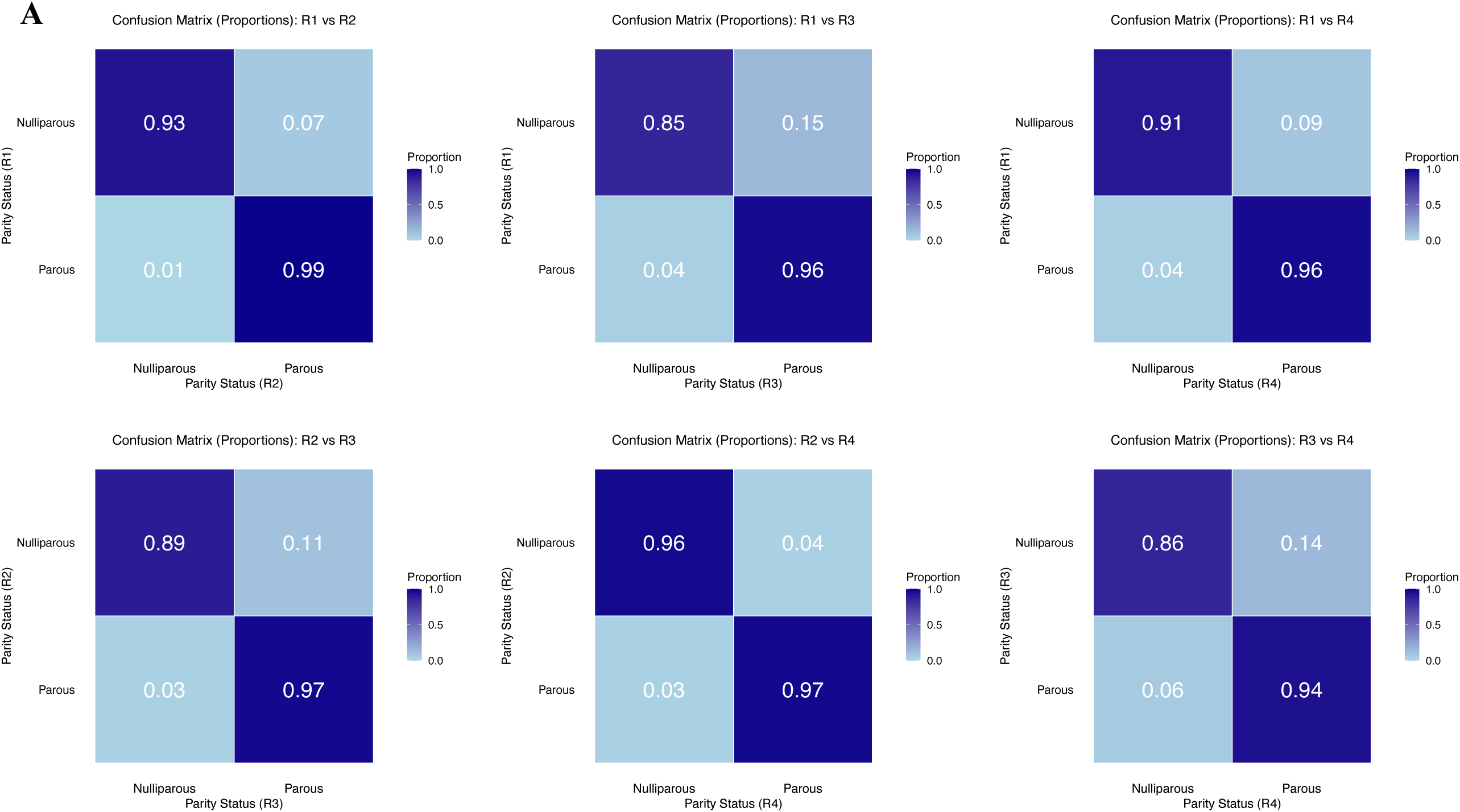
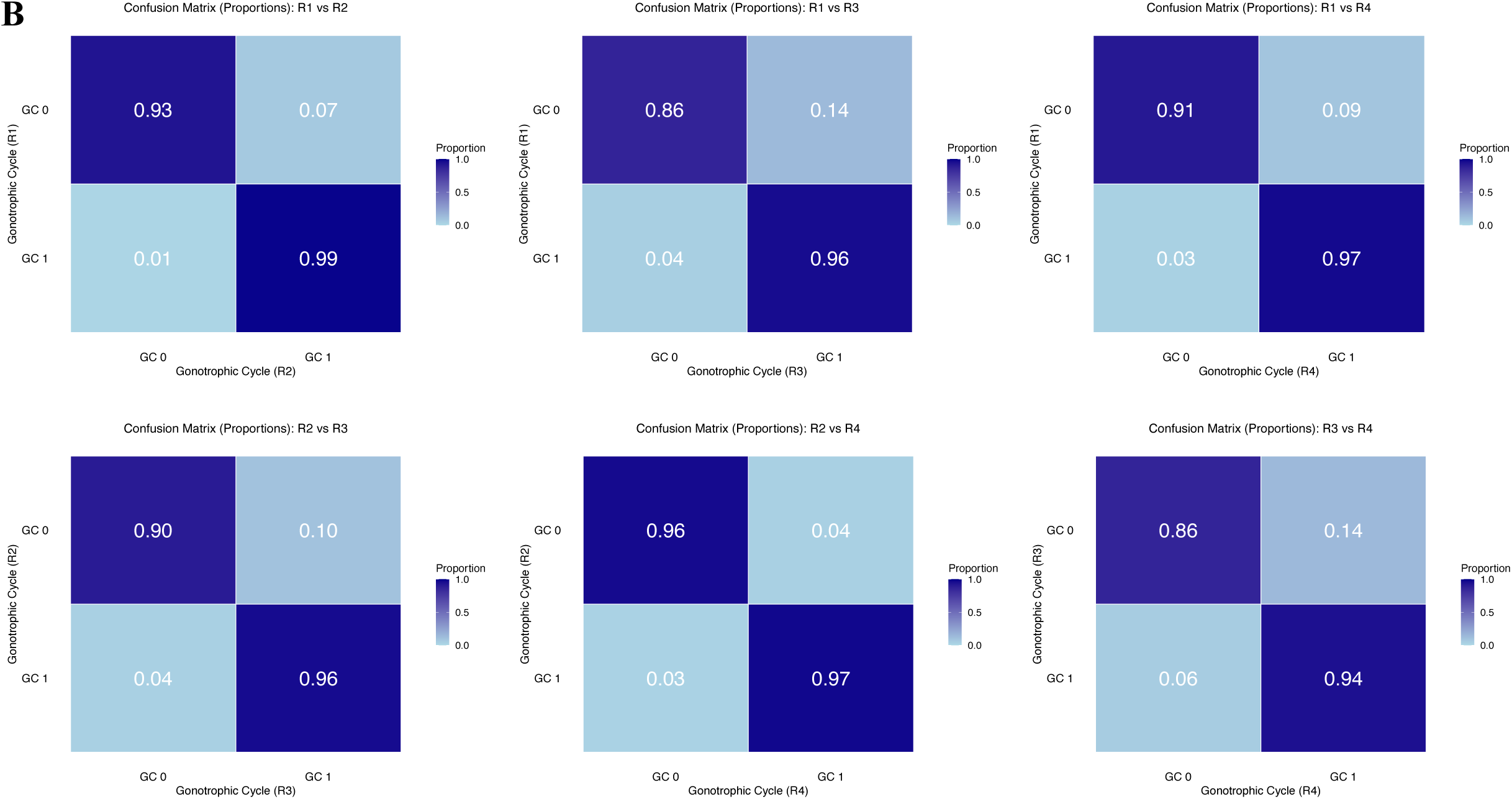
Precision of A) parity and B) gonotrophic cycle classification in field-collected *An. funestus*. Confusion matrices showing pairwise agreement in parity (nulliparous vs parous) and gonotrophic cycle classification (GC 0 and GC 1) between researchers (R1 vs R2, R1 vs R3, R1 vs R4, R2 vs R3, R2 vs R4, and R3 vs R4).

## Discussion

This study provides a comprehensive evaluation of the accuracy and precision of the Polovodova method for parity and gonotrophic cycle classification in two major African malaria vectors, *An. arabiensis* and *An. funestus*, under both controlled laboratory and field conditions. Overall, we found that the Polovodova method was both accurate and precise for distinguishing nulliparous from parous mosquitoes and for identifying those that had undergone one or two gonotrophic cycles, with high inter-rater reliability across researchers for both species. However, the method performed substantially less well at classifying mosquitoes into specific gonotrophic cycle categories, with both accuracy and precision showing a pronounced and progressive decline at later gonotrophic cycles (GC2 and above). Systematic underestimation of older age classes was evident across both species and all independent age graders, with accuracy declining from approximately 84-97% at GC1 to 33-46% at GC4. Despite these limitations in GC classification, the high reliability of the method for parity assessment supports its continued use in laboratory and field surveillance. The constraints identified for gonotrophic cycle level classification have important implications for reliable assessment of mosquito age structure in the field, particularly with respect to the proportion surviving to older age categories as required for transmission intensity estimation and the evaluation of vector control interventions.

The high accuracy of parity classification observed here is broadly consistent with findings from previous evaluations of ovarian dissection methods in *Anopheles* mosquitoes. For instance, Pretorius et al. (Pretorius et al. 2023), who validated parity assessment using the ovarian tracheation technique in *An. gambiae s.l.*, reported similarly high inter-rater reliability for both freshly killed insectary-reared mosquitoes and field-caught mosquitoes (Pretorius et al. 2023). In contrast, Hugo et al. (Hugo et al. 2008) evaluated the Polovodova method in *Aedes vigilax* and *Culex annulirostris* and reported markedly lower overall classification accuracy of 57.5%, reflecting the greater difficulty and poorer performance of this method for those species (Hugo et al. 2008, Johnson et al. 2020). The substantially higher agreement observed in the present study may reflect the more distinct ovarian morphological changes associated with oviposition in African *Anopheles* mosquitoes (Detinova 1945, 1962, Gillies and Wilkes 1965).

Our findings for gonotrophic cycle (GC) classification are consistent with previous evaluations of the Polovodova method, although direct comparisons are limited by differences in mosquito species, study design, and the number of GC categories assessed. Anagonou *et al*. (Anagonou et al. 2015) evaluated the Polovodova method in *An. gambiae s.s.* in Benin using mosquitoes of known egg-laying histories and found that while this technique was effective for distinguishing nulliparous and uniparous mosquitoes it performed poorly in multiparous females, with the proportion of ovarioles displaying the diagnostic number of dilatations (*i.e.* the number of dilatations exactly matching the number of completed gonotrophic cycles) declining progressively from approximately one third at GC1, to one tenth at GC2, and one twentieth at GC3 (Anagonou et al. 2015). A similar trend was observed in our study, where agreement with the assumed reference dataset declined from GC1 to GC4. Comparable observations have also been reported for *Culex annulirostris* (Hugo et al. 2008), suggesting that this decline is not species-specific but reflects reflects a fundamental limitation of the method: the progressive reduction in the proportion of ovarioles displaying the expected number of dilatations with increasing gonotrophic cycles (Hoc and Wilkes 1995, Hugo et al. 2008).

Analysis of misclassification patterns showed that parity errors were uncommon and broadly balanced across researchers and species. Parous females were slightly more often misclassified as nulliparous (4-6%) than nulliparous females as parous (2-3%), but errors in both directions were small, indicating little systematic bias in estimated parity rates. In contrast, gonotrophic-cycle misclassification was strongly directional: when errors occurred, mosquitoes were almost invariably assigned one cycle below the assumed reference category, such as GC4 being classified as GC3. This underestimation increased with age, with agreement declining to 33-46% at GC4 in both species. These findings suggest that the Polovodova method becomes progressively less reliable at later gonotrophic cycles, likely because repeated ovarian dilatations become increasingly difficult to distinguish. Its performance may therefore depend strongly on reader training and experience, while also reflecting an inherent practical limit of the technique for resolving older mosquitoes.

This systematic bias has important epidemiological implications. Underestimating later gonotrophic cycles would make mosquito populations appear younger and could consequently underestimate transmission potential, because only females surviving beyond the parasite’s extrinsic incubation period can transmit malaria (Detinova 1945, 1962, Beklemishev et al. 1959, Gillies and Wilkes 1965). The consequences may be limited when evaluating rapidly acting interventions, such as effective pyrethroid treated nets, that sharply reduce mosquito survival. However, for delayed action interventions such as chlorfenapyr containing Interceptor G2 nets (N’Guessan et al. 2016, Gervas et al. 2026), failure to detect reductions in mosquitoes reaching later gonotrophic cycles could underestimate their impact on longevity and transmission. One approach to addressing this limitation is to apply correction factors derived from laboratory validation studies to adjust observed gonotrophic cycle distributions.

Several biological and technical factors likely contributed to the declining performance of GC classification at later gonotrophic cycles. Biologically, older dilatations may adhere to the calyx wall or become obscured by structural changes in ovarian tissue (Hoc and Wilkes 1995) and increased pigment deposition associated with repeated gonotrophic cycling, reducing the contrast between dilatations and surrounding structures and complicating interpretation (Hugo et al. 2008, Anagonou et al. 2015). Technically, prior handling of ovaries by another researcher, accidental cutting of dilatations during dissection, and obscuring of ovarian structures by residual blood meals or sugar deposits on slides were identified as additional sources of error. Field-collected mosquitoes were perceived as somewhat easier to dissect than laboratory-reared ones, possibly reflecting differences in tissue condition between wild and insectary-reared mosquitoes. These combined biological and technical challenges reinforce the need for standardised training protocols and careful slide preparation to maximise reproducibility of the Polovodova method across observers (Hugo et al. 2008, Pretorius et al. 2023).

Precision of both parity and GC classification in field-collected mosquitoes was high and broadly consistent with laboratory findings. Parous females with visible ovarian sac were identified exclusively among field-collected samples, reflecting natural variation in gonotrophic cycle length between individuals rather than random sampling across cycle stages mosquitoes captured in this transitional state had completed oviposition and resumed host-seeking earlier than average, consistent with known variation in cycle duration in Afrotropical malaria vectors (Charlwood et al. 2018, 2023). Classification was further complicated by follicular sacs in varying states of closure within the same mosquito, making distinction between parous females with visible sac and completed GC1 particularly difficult in field samples. While the small number of parous females with sac identified in field collections (n < 20 per species) precluded formal precision analysis of this category, their presence highlights an important source of potential misclassification if not clearly distinguished from GC1. These challenges are compounded by the inherent subjectivity of morphological scoring (Gillies and Wilkes 1965, Hugo et al. 2008).

Additionally, *An. funestus* consistently showed slightly lower agreement than *An. arabiensis* across both parity and GC classification, in terms of both accuracy against the R0 reference and inter-rater precision. To our knowledge, no previous study has specifically compared Polovodova method performance between these two species. However, species-level differences in dissection difficulty have been noted in the literature, with Gillies and Wilkes (Gillies and Wilkes 1965) reported greater difficulty applying the Polovodova method to *An. funestus* than *An. gambiae* due to its smaller body size and more delicate ovarian structures, and Charlwood et al. (Charlwood et al. 1985) similarly noted that *An. farauti* was easier to dissect than *An. gambiae*. Consistent with this, graders in the present study perceived *An. funestus* as considerably more difficult to dissect than *An. arabiensis*, attributing this to the same morphological characteristics. Both species are also known to exhibit gonotrophic discordance under certain circumstances, whereby females may require more than one blood meal to complete a single gonotrophic cycle (Briegel and Hörler 1993, Beier 1996), which may introduce additional biological variability in GC classification.

It should be noted that expert dissectors with extensive field experience have reported accurate gonotrophic cycle classification up to six gonotrophic cycles in field-caught mosquitoes, indicating that method performance is strongly influenced by dissector skill and experience (Charlwood et al. 1985). The error rates quantified here therefore likely reflect performance at a defined training level rather than an absolute limitation of the method itself. Charlwood et al. also emphasised the importance of examining the entire ovary for accurate age estimation, consistent with our observation that the proportion of ovarioles displaying the correct number of dilatations decreases progressively with increasing gonotrophic cycles (Charlwood et al. 1985). All researchers in the present study examined ovarioles from both ovaries for each mosquito; despite this, classification accuracy still declined substantially at later cycles, suggesting that the observed underestimation primarily reflects the fundamental biological reduction in the proportion of ovarioles retaining a complete dilatation record with successive gonotrophic cycling rather than incomplete ovarian examination. Nevertheless, it is likely that researchers with greater dissection experience than those involved in the present study would achieve improved classification accuracy at later gonotrophic cycles, and future studies should consider recruiting expert-level dissectors alongside less experienced researchers to better characterise the relative contributions of biological and technical factors to GC misclassification.

Several limitations of this study warrant acknowledgement. A confirmed individual-level ground truth (reference) for GC classification was not available in this study. While direct observation of oviposition provided a reliable ground truth for parity status and GC1 classifications after the first blood meal, the GC status of mosquitoes at later blood-feeding events had to be estimated using an assumed reference dataset derived from blood-feeding histories and oviposition records. Although the assumptions underlying this model were biologically informed and reasonable, the use of the derived reference introduces uncertainty that a true individual-level ground truth would not. If the true GC of some mosquitoes was overestimated (e.g. by assuming a lower rate of gonotrophic discordance than occurred in these cohorts) this may have generated a systematic bias unrelated to any limitation of the Polovodova method itself. While we believe this source of error is unlikely to fully explain the magnitude and consistency of GC underestimation, we cannot rule it out as a potential limitation of our study design. Future studies tracking individual oviposition histories across all gonotrophic cycles would provide a stronger foundation for evaluating GC classification accuracy. A further limitation was the inability to evaluate precision beyond GC1 for field-collected mosquitoes, as field populations predominantly comprised nulliparous and GC1 individuals, with very few higher-cycle mosquitoes available for analysis, consistent with natural field population age structures where older mosquitoes are typically underrepresented due to mortality and ecological factors (Gillies and Wilkes 1965). Given that laboratory results demonstrated declining precision with increasing GC number, this represents an important gap for future field validation studies.

These findings have direct practical implications for national malaria vector surveillance programmes. These findings have direct practical implications for national malaria vector surveillance programmes. The Polovodova method can be reliably used for parity classification and early GC assessment (GC1-GC2), while GC3-GC4 classifications should be interpreted cautiously. To improve reliability, programmes should consider implementing double or consensus reading workflows, standardised training protocols, and clear operating procedures are essential for consistent application with particular attention needed for *An. funestus*. Integration with complementary approaches such as infrared spectroscopy (Mayagaya et al. 2009, Sikulu-Lord et al. 2018, Siria et al. 2022) and transcriptional biomarkers (Cook et al. 2007, Tuwei et al. 2025) is recommended where resources permit.

Overall, the Polovodova method remains a robust and reproducible tool for parity classification in African malaria vectors across laboratory and field settings. However, its utility for detailed gonotrophic cycle estimation is constrained by progressive and systematic underestimation at later GC stages. Future work should investigate whether correction factors derived from laboratory validation studies can improve the reliability of field-based GC estimates and explore integration of the Polovodova method with complementary approaches to improve the resolution and reproducibility in large-scale surveillance programmes.

### Conclusion

This study provides the most comprehensive evaluation to date of the accuracy and precision of the Polovodova method for parity and gonotrophic cycle classification in two major African malaria vectors, *An. arabiensis* and *An. funestus*, under both controlled laboratory and field settings. The method demonstrated high accuracy and almost perfect inter-rater reliability for parity classification across both species and settings, supporting its continued use as a robust tool for broad reproductive status assessment in mosquito surveillance programmes. However, its utility for detailed gonotrophic cycle estimation is substantially constrained by progressive and systematic underestimation of older age classes a limitation with important implications for the accuracy of malaria transmission intensity estimates and the evaluation of vector control interventions, particularly those with delayed modes of action. Field-based evaluation confirmed that the method can be applied consistently by multiple researchers under natural conditions, though the restricted age structure of field populations limited assessment of precision beyond GC1. To maximise reliability in operational settings, national malaria control programmes should implement consensus or double-reading workflows for late-cycle specimens, where disagreement is most likely and the consequences of misclassification are greatest. Standardised training protocols and clear operating procedures are essential prerequisites for consistent application across researchers, with particular attention warranted for *An. funestus*, which was consistently perceived as more challenging to dissect and showed slightly lower inter-rater agreement than *An. arabiensis*. Where resources permit, integration of the Polovodova method with complementary spectroscopic approaches such as near- and mid-infrared spectroscopy is recommended to strengthen age-grading resolution beyond what dissection-based methods can reliably achieve alone.

## Acknowledgements

We would like to express our gratitude to all study participants for their time and valuable insights into the whole process of this research. We thank Jacques Derek Charlwood for mosquito identification and dissection trainings provided prior the start of the project. We thank Paul Johnson for support with statistical analysis. We also thank the technician team: Agapiti Soka, Peter Mchele, and Ramadhani Saidi for their technical assistance with mosquito collections, dissections, and age-grading. We additionally thank Rukiyah Mohammed for administrative support throughout the project.

## Abbreviations

EIP: Extrinsic incubation period
GC: Gonotrophic cycle
P1: Parous 1
P2: Parous 2
P3: Parous 3
P4: Parous 4
MALDITOF-MS: Matrix-assisted laser desorption ionization time-of-flight mass spectrometry
WHO: World Health Organisation
MIRS: Mid-infrared spectroscopy
NIRS: Near-infrared spectroscopy

## Authors contributions

DJS was the lead study investigator who was involved in the study design, data collection, data entry and analysis, interpretation of the results, and writing of the manuscript. HSN assisted in study design and analysis. GT and LM were involved in data collection. HSN, EH and MJ were involved in the study design and in manuscript writing (review and editing). All authors read and approved the final manuscript. FO, FB, and HMF were involved in the study design, supervision, and critical revision of the manuscript.

## Availability of data and materials

All the data for this study is available upon reasonable request.

## Ethics approval

Ethical consideration was sought prior to the work; ethical approval was sought from the Ifakara Health institutional review board (IRB), reference number (IHI/IRB/EXT/No: 02-2025), and National Institute for Medical Research reference number (NIMR/HQ/R.8a/Vol. IX/4112). Permission to publish was sought from the National Institute for Medical Research (NIMR) with a reference number Ref No. BD.242/437/01C/231.

