## Supplementary material for "How reliable is the use of ovarian dilatations for mosquito age-grading? A blinded multi-rater validation of the Polovodova method in *Anopheles* mosquitoes": Supplementary Material_SIRIA.pdf

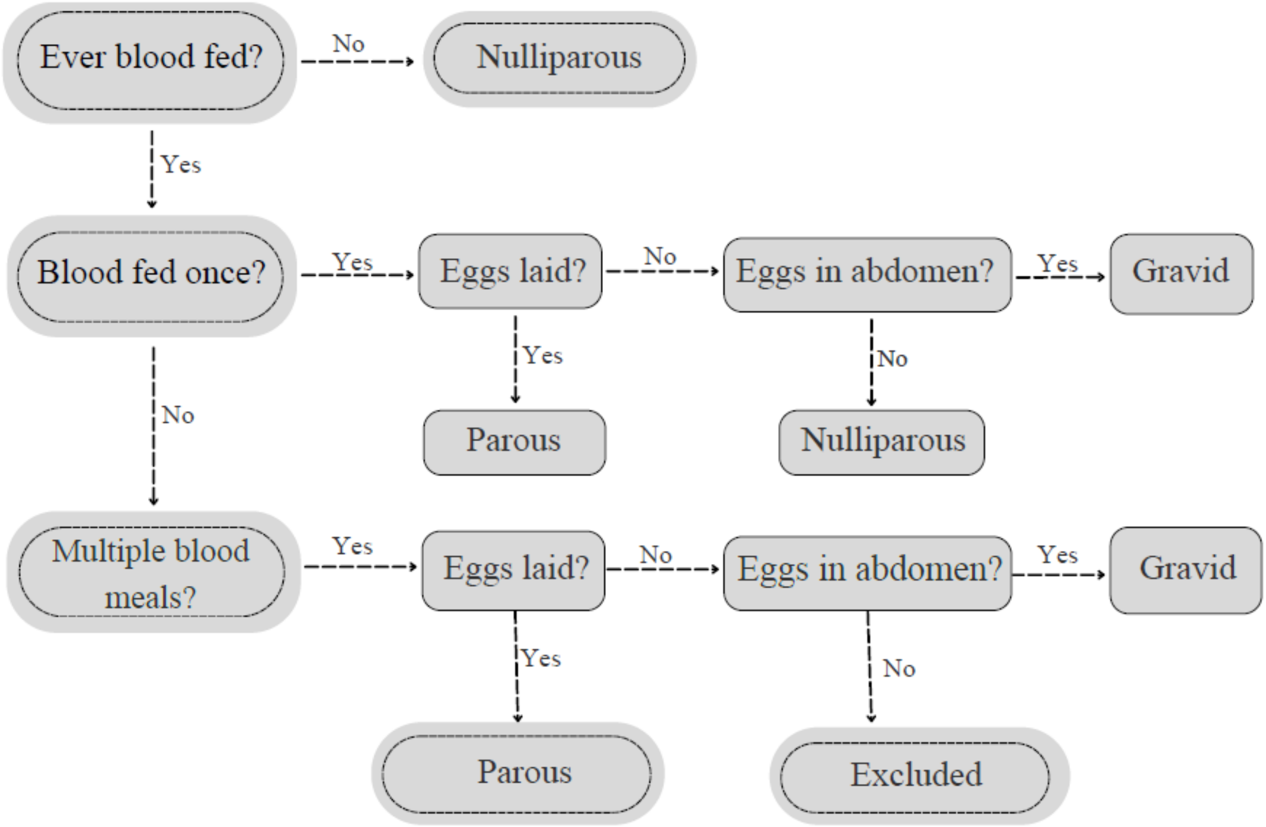

**Figure S1. Establishment of parity 'reference group'**. Flow diagram illustrating the true classification of parous and nulliparous mosquitoes based on observations on blood-feeding history and oviposition outcome (based on R0), and gravid status (based on dissections made by R1-R3). Mosquitoes were classified as nulliparous, parous, or gravid. Individuals lacking clear parity determination were excluded.

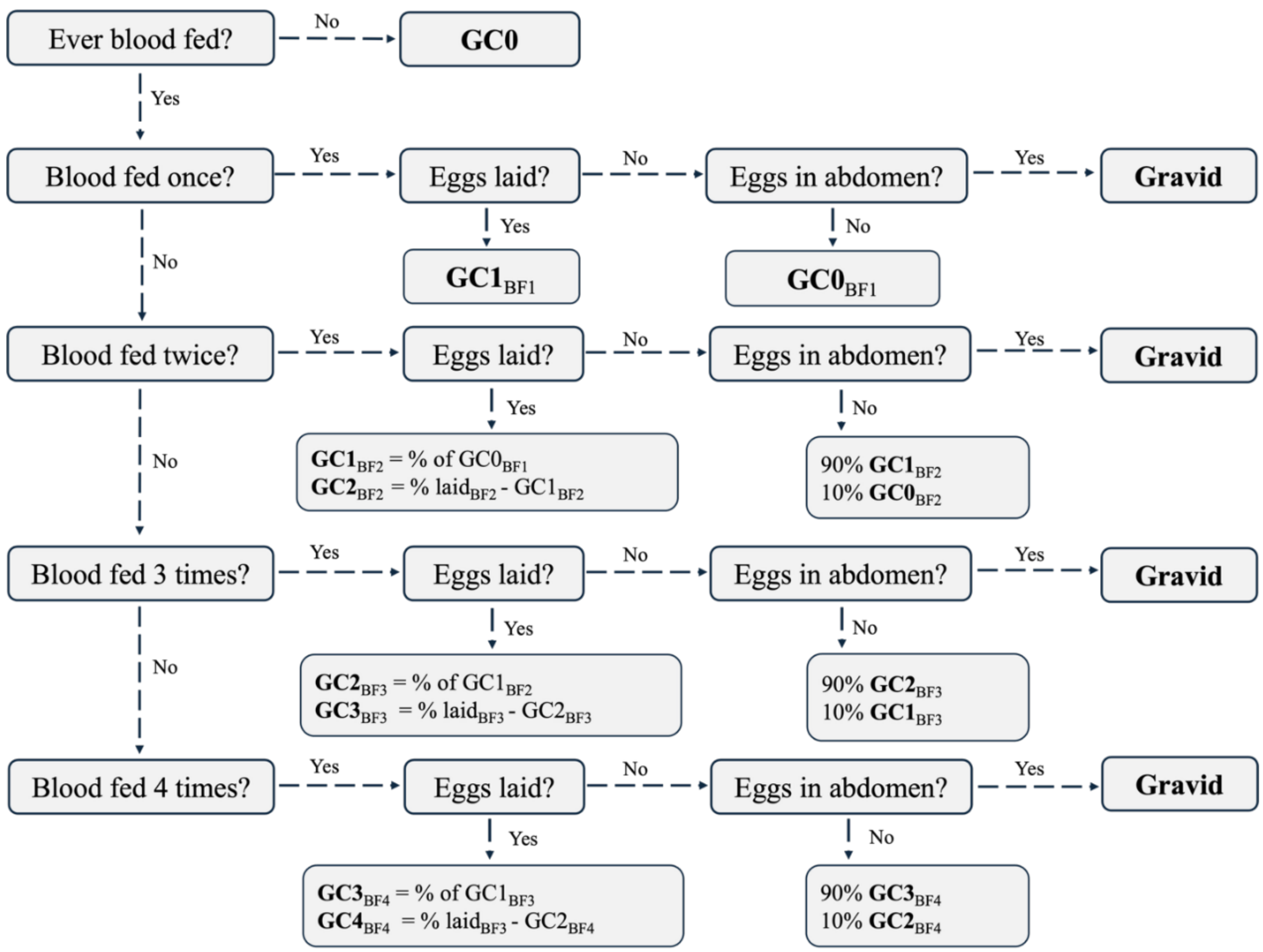

**Figure S2**. **Establishment of assumed gonotrophic cycles.** Schematic showing the decision framework used to infer assumed gonotrophic cycle (AR) categories based on blood-feeding history, oviposition outcome, and gravid status. Mosquitoes were classified as nulliparous, parous (GC1-GC4), or gravid using biologically informed rules applied iteratively across successive blood meals.

**Table S1.** Inter-rater Agreement between reference data/ground truth (R0) and each of the other rates (R1, R2, R3)

| Species | Researcher 0 | | Researcher 1 | | IRR-*Kappa* | P-Value |
|---|---|---|---|---|---|---|
| | Nulliparous n/N (prop) | Parous n/N (prop) | Nulliparous n/N (prop) | Parous n/N (prop) | | |
| *An. arabiensis* | 439/472 (0.93) | 1254/1332 (0.94) | 439/517 (0.85) | 1254/1263 (0.99) | 0.87 [0.83, 0.92] | <0.001 |
| | | | Researcher 2 | | | |
| | 438/472 (0.93) | 1262/1332 (0.95) | 438/508 (0.86) | 1262/1272 (0.99) | 0.88 [0.83, 0.92] | <0.001 |
| | | | Researcher 3 | | | |
| | 440/472 (0.93) | 1265/1332 (0.95) | 440/507 (0.87) | 1265/1273 (0.99) | 0.89 [0.84, 0.93] | <0.001 |
| | Researcher 0 | | Researcher 1 | | | |
| *An. funestus* | 362/365 (0.99) | 1198/1279 (0.94) | 362/442 (0.82) | 1198/1199 (1) | 0.86 [0.81, 0.91] | < 0.001 |
| | | | Researcher 2 | | | |
| | 362/365 (0.99) | 1217/1279 (0.95) | 362/423 (0.86) | 1217/1218 (1) | 0.89 [0.84, 0.94] | < 0.001 |
| | | | Researcher 3 | | | |
| | 363/365 (0.99) | 1233/1279 (0.96) | 363/408 (0.89) | 1233/1233 (1) | 0.92 [0.88, 0.97] | < 0.001 |

**Table S2** Agreement (%) between researcher (R1-R3) gonotrophic cycle classifications and the AR dataset for both *An. arabiensis* and *An. funestus*

| ***An. arabiensis*** | | | | | |
|---|---|---|---|---|---|
| **Blood-feeding** | **R1 (%)** | **R2 (%)** | **R3 (%)** | **Mean (%)** | **SD** |
| 0 BF | 100 | 100 | 100 | 100 | 0.0 |
| 1 BF | 84 | 86 | 86 | 86 | 1.1 |
| 2 BF | 88 | 88 | 97 | 91 | 5.5 |
| 3 BF | 74 | 76 | 76 | 75 | 1.0 |
| 4 BF | 45 | 46 | 44 | 45 | 0.8 |
| ***An. funestus*** | | | | | |
| 0 BF | 100 | 100 | 100 | 100 | 0.0 |
| 1 BF | 79 | 86 | 88 | 84 | 5.1 |
| 2 BF | 86 | 94 | 97 | 93 | 5.6 |
| 3 BF | 69 | 69 | 65 | 68 | 2.2 |
| 4 BF | 33 | 36 | 35 | 35 | 1.3 |

**Table S3.** Multinomial-Dirichlet overlap scores for *An. arabiensis*

0 = no overlap with AR | 1 = perfect match |

| Blood Feeding | Comparison | Score | 95% CI |
|---|---|---|---|
| BF0 | AR vs R1 | 1.00 | [0.99-0.99] |
| BF0 | AR vs R2 | 1.00 | [0.99-0.99] |
| BF0 | AR vs R3 | 1.00 | [0.99-0.99] |
| BF1 | AR vs R1 | 0.84 | [0.80-0.89] |
| BF1 | AR vs R2 | 0.86 | [0.82-0.90] |
| BF1 | AR vs R3 | 0.86 | [0.82-0.90] |
| BF2 | AR vs R1 | 0.88 | [0.83-0.92] |
| BF2 | AR vs R2 | 0.88 | [0.83-0.92] |
| BF2 | AR vs R3 | 0.97 | [0.94-0.99] |
| BF3 | AR vs R1 | 0.75 | [0.69-0.79] |
| BF3 | AR vs R2 | 0.76 | [0.71-0.81] |
| BF3 | AR vs R3 | 0.76 | [0.71-0.81] |
| BF4 | AR vs R1 | 0.46 | [0.43-0.48] |
| BF4 | AR vs R2 | 0.46 | [0.43–0.49] |
| BF4 | AR vs R3 | 0.45 | [0.42-0.48] |

Excellent (⩾0.90) | Good/Moderate (0.70-0.89) | Poor (<0.70)

**Table S4.** Multinomial-Dirichlet overlap scores for *An. funestus*

0 = no overlap with AR | 1 = perfect match |

| Blood Feeding | Comparison | Score | 95% CI |
|---|---|---|---|
| BF0 | AR vs R1 | 1.00 | [0.99-0.99] |
| BF0 | AR vs R2 | 1.00 | [0.99-0.99] |
| BF0 | AR vs R3 | 1.00 | [0.99-0.99] |
| BF1 | AR vs R1 | 0.79 | [0.73-0.83] |
| BF1 | AR vs R2 | 0.86 | [0.81-0.90] |
| BF1 | AR vs R3 | 0.88 | [0.84-0.92] |
| BF2 | AR vs R1 | 0.87 | [0.81-0.92] |
| BF2 | AR vs R2 | 0.95 | [0.89-0.99] |
| BF2 | AR vs R3 | 0.97 | [0.92-0.99] |
| BF3 | AR vs R1 | 0.69 | [0.64-0.74] |
| BF3 | AR vs R2 | 0.69 | [0.64-0.75] |
| BF3 | AR vs R3 | 0.66 | [0.65-0.71] |
| BF4 | AR vs R1 | 0.34 | [0.32-0.36] |
| BF4 | AR vs R2 | 0.36 | [0.33-0.39] |
| BF4 | AR vs R3 | 0.36 | [0.33-0.39] |

Excellent (⩾0.90) | Good/Moderate (0.70-0.89) | Poor (<0.70)

**Table S5.** Inter-rater reliability of parity classification (nulliparous vs parous) by three researchers for *An. arabiensis* and *An. funestus*

| **Species** | **Researcher** | | **Researcher** | | **Cohen's IRR *Kappa*(κ) [95 % CI]** | **P-value** |
|---|---|---|---|---|---|---|
| | Nulliparous n/N (prop) | Parous n/N (prop) | Nulliparous n/N (prop) | Parous n/N (prop) | | |
| | **Researcher 1** | | **Researcher 2** | | | |
| | 446/531 (0.84) | 1396/1477 (0.95) | 446/518 (0.86) | 1396/1490 (0.94) | 0.80 [0.76, 0.84] | <0.001 |
| | **Researcher 1** | | **Researcher 3** | | | |
| ***An. arabiensis*** | 445/531 (0.84) | 1401/1477 (0.95) | 445/520 (0.86) | 1401/1499 (0.93) | 0.80 [0.77, 0.84] | <0.001 |
| | **Researcher 2** | | **Researcher 3** | | | |
| | 444/518 (0.86) | 1415/1490 (0.95) | 444/520 (0.85) | 1415/1499 (0.94) | 0.82 [0.78, 0.85] | <0.001 |
| | **Researcher 1** | | **Researcher 2** | | | |
| ***An. funestus*** | 380/466 (0.82) | 1179/1244 (0.95) | 380/445 (0.85) | 1179/1265 (0.93) | 0.80 [0.76, 0.84] | < 0.001 |
| | **Researcher 1** | | **Researcher 3** | | | |
| | 387/466 (0.83) | 1196/1244 (0.96) | 387/435 (0.89) | 1196/1275 (0.94) | 0.83 [0.79, 0.87] | < 0.001 |
| | **Researcher 2** | | **Researcher 3** | | | |
| | 395/445 (0.89) | 1225/1265 (0.97) | 395/435 (0.91) | 1225/1275 (0.96) | 0.88 [0.84, 0.92] | < 0.001 |

**Table S6.** Pairwise inter-rater agreement in gonotrophic cycle classification (GC0-GC3) between researchers for *Anopheles arabiensis* and *Anopheles funestus*.

| Species | Researcher | | | | Researcher | | | | Cohen's IRR *Kappa*(κ) [95 % CI] | P-value |
|---|---|---|---|---|---|---|---|---|---|---|
| | GC0 n/N (prop) | GC1 n/N (prop) | GC2 n/N (prop) | GC3 n/N (prop) | GC0 n/N (prop) | GC1 n/N (prop) | GC2 n/N (prop) | GC3 n/N (prop) | | |
| *Anopheles arabiensis* | Researcher 1 | | | | Researcher 2 | | | | | |
| | 446/531 (0.84) | 286/440 (0.65) | 316/528 (0.60) | 394/516 (0.76) | 446/518 (0.86) | 286/456 (0.63) | 316/521 (0.61) | 394/520 (0.76) | 0.58 [0.56, 0.61] | <0.001 |
| | Researcher 1 | | | | Researcher 3 | | | | | |
| | 445/531 (0.84) | 265/440 (0.60) | 354/528 (0.67) | 393/516 (0.76) | 445/520 (0.86) | 265/408 (0.65) | 354/579 (0.61) | 393/508 (0.77) | 0.60 [0.57, 0.63] | <0.001 |
| | Researcher 2 | | | | Researcher 3 | | | | | |
| | 444/518 (0.86) | 269/456 (0.59) | 338/521 (0.65) | 391/520 (0.75) | 444/520 (0.85) | 269/408 (0.66) | 338/579 (0.58) | 391/508 (0.77) | 0.59 [0.57, 0.63] | <0.001 |
| *Anopheles funestus* | Researcher 1 | | | | Researcher 2 | | | | | |
| | 380/465 (0.82) | 209/356 (0.59) | 286/479 (0.60) | 279/406 (0.69) | 380/443 (0.86) | 209/362 (0.58) | 286/498 (0.57) | 279/403 (0.69) | 0.55 [0.52, 0.57] | < 0.001 |
| | Researcher 1 | | | | Researcher 3 | | | | | |
| | 387/466 (0.83) | 224/358 (0.63) | 296/479 (0.62) | 272/405 (0.67) | 387/435 (0.89) | 224/366 (0.61) | 296/517 (0.57) | 272/390 (0.7) | 0.57 [0.54, 0.59] | < 0.001 |
| | Researcher 2 | | | | Researcher 3 | | | | | |
| | 393/443 (0.89) | 249/363 (0.69) | 302/497 (0.61) | 261/403 (0.65) | 393/432 (0.91) | 249/367 (0.68) | 302/517 (0.58) | 261/390 (0.67) | 0.58 [0.56, 0.61] | < 0.001 |

**Table S7.** Inter-rater reliability for parity classification (Field-collected mosquitoes)

| Species | Overall (Fleiss' κ) | Pairwise Comparison | Cohen's κ | 95% CI | p-value |
|---|---|---|---|---|---|
| *An. arabiensis* (n=559) | **0.87** | R1 vs R2 | 0.933 | [0.89-0.97] | <0.001 |
| | | R1 vs R3 | 0.845 | [0.79-0.89] | <0.001 |
| | | R1 vs R4 | 0.865 | [0.81-0.92] | <0.001 |
| | | R2 vs R3 | 0.880 | [0.83-0.93] | <0.001 |
| | | R2 vs R4 | 0.901 | [0.86-0.95] | <0.001 |
| | | R3 vs R4 | 0.822 | [0.76-0.88] | <0.001 |
| *An. funestus* (n=507) | **0.86** | R1 vs R2 | 0.927 | [0.89-0.97] | <0.001 |
| | | R1 vs R3 | 0.821 | [0.76-0.88] | <0.001 |
| | | R1 vs R4 | 0.863 | [0.81-0.92] | <0.001 |
| | | R2 vs R3 | 0.859 | [0.81-0.91] | <0.001 |
| | | R2 vs R4 | 0.912 | [0.87-0.95] | <0.001 |
| | | R3 vs R4 | 0.785 | [0.72-0.85] | <0.001 |

**Table S8.** Inter-rater reliability for GC classification (Field-collected mosquitoes)

| **Species** | **Overall (Fleiss' κ)** | **Pairwise Comparison** | **Cohen's κ** | **95% CI** | **p-value** |
|---|---|---|---|---|---|
| *An. arabiensis (n=521)* | **0.88** | R1 vs R2 | 0.93 | [0.89-0.97] | <0.001 |
| | | R1 vs R3 | 0.85 | [0.79-0.90] | <0.001 |
| | | R1 vs R4 | 0.88 | [0.83-0.93] | <0.001 |
| | | R2 vs R3 | 0.88 | [0.83-0.93] | <0.001 |
| | | R2 vs R4 | 0.91 | [0.87-0.96] | <0.001 |
| | | R3 vs R4 | 0.83 | [0.77-0.89] | <0.001 |
| *An. funestus* (n=459) | **0.87** | R1 vs R2 | 0.93 | [0.89-0.97] | <0.001 |
| | | R1 vs R3 | 0.83 | [0.77-0.87] | <0.001 |
| | | R1 vs R4 | 0.88 | [0.83-0.93] | <0.001 |
| | | R2 vs R3 | 0.86 | [0.80-0.91] | <0.001 |
| | | R2 vs R4 | 0.92 | [0.88-0.97] | <0.001 |
| | | R3 vs R4 | 0.79 | [0.73-0.86] | <0.001 |
